# Homeostatic scaling is driven by a translation-dependent degradation axis that recruits miRISC remodeling

**DOI:** 10.1101/2020.04.01.020164

**Authors:** Balakumar Srinivasan, Sarbani Samaddar, Sivaram V.S. Mylavarapu, James P. Clement, Sourav Banerjee

**Affiliations:** National Brain Research Centre, NH-8. Nainwal Mode, Manesar-122052, Haryana, India; Regional Centre for Biotechnology, NCR-Biotech Science Cluster, Faridabad-Gurgaon Expressway, Faridabad-121001, Haryana, India; Neuroscience Unit, Jawaharlal Nehru Centre for Avanced Scientific Research, Jakkur, Bengaluru-560064, Karnataka, India

**Author notes:** These authors contributed equally to this work (Balakumar Srinivasan, Sarbani Samaddar). Correspondence OR (S.B).

## Abstract

Homeostatic scaling in neurons has been attributed to the individual contribution of either translation or degradation; however there remains limited insight towards understanding how the interplay between the two processes effectuates synaptic homeostasis. Here, we report that a co-dependence between protein synthesis and degradation mechanisms drives synaptic homeostasis whereas abrogation of either prevents it. Coordination between the two processes is achieved through the formation of a tripartite complex between translation regulators, the 26S proteasome and the miRNA-induced-silencing-complex (miRISC) components such as Argonaute, MOV10 and Trim32 on actively translating transcripts or polysomes. The components of this ternary complex directly interact with each other in an RNA-dependent manner. Disruption of polysomes abolishes this ternary interaction, suggesting that translating RNAs facilitate the combinatorial action of the proteasome and the translational apparatus. We identify that synaptic downscaling involves miRISC remodeling which entails the mTORC1-dependent translation of Trim32, an E3 ligase and the subsequent degradation of its target, MOV10 *via* the phosphorylation of p70 S6 kinase. We find that the E3 ligase Trim32 specifically polyubiquitinates MOV10 for its degradation during synaptic downscaling. MOV10 degradation alone is sufficient to invoke downscaling by enhancing Arc translation through its 3’ UTR and causing the subsequent removal of post-synaptic AMPA receptors. Synaptic scaling was occluded when we depleted Trim32 and overexpressed MOV10 in neurons, suggesting that the Trim32-MOV10 axis is necessary for synaptic downscaling. We propose a mechanism that exploits a translation-driven protein degradation paradigm to invoke miRISC remodeling and induce homeostatic scaling during chronic network activity.

## Introduction

Neurons employ a unique strategy, known as synaptic scaling, to counter the run-away excitation and subsequent loss of input specificity that arise due to Hebbian changes; they rely on a compensatory remodeling of synapses throughout the network while maintaining differences in their synaptic weightage [1–6]. Synaptic scaling is achieved by a complex interplay of sensors and effectors within neurons that serve to oppose global fluctuations in a network and establish synaptic homeostasis by modifying post-synaptic glutamatergic currents in a cell-autonomous manner [7–9]. In the context of homeostatic scaling, ‘sensors’ are classified as molecules that sense deviations in the overall network activity and ‘effectors’ scale the neuronal output commensurately.

Till date not much is known about the repertoire of molecular “sensor” cascades that serve to link events where neurons sense deviations in the network firing rate and subsequently initiate the scaling process. Few molecular sensors have been identified; the eukaryotic elongation factor eEF2 and its dedicated kinase, eEF2 kinase or CamKIII are the two reported thus far [10]. One cascade discovered in this context is the mTORC1 (mammalian Target Of Rapamycin Complex-1) signalling pathway that regulates presynaptic compensation in neurons by promoting BDNF synthesis in the post-synaptic compartment [11,12]. In contrast, AMPA-receptors (AMPARs) have been identified, by overwhelming consensus, to be the predominant “end-point-effectors” in all paradigms of synaptic scaling [13–16]. Unlike NMDARs, AMPARs undergo *de novo* translation during network destabilizations [17] and chronic changes in the post-synaptic response during scaling has been attributed to the abundance of surface AMPARs (GluA1 and GluA2 subunits) [18]. Among the key modifiers of AMPAR expression, miRNAs are known to play pivotal roles in synaptic scaling [19–22]. Relief from translational repression by miRNAs necessitates that mRNAs exit the functional miRISC (microRNA induced silencing complex). This requires miRISC to undergo dynamic changes in its composition [23,24], a cellular phenomenon previously termed as miRISC remodeling [25]. However, what remains surprising is our lack of knowledge about how compositional changes within the miRISC are achieved during scaling.

The requirement for discrete sets of sensors and effectors is fulfilled within neurons through varied mechanisms including translation and ubiquitin-mediated proteasomal (UPS) degradation. An enhanced degradation of post-synaptic-density (PSD) proteins including GluA1 and GluA2 has been observed in contexts of altered network excitability [26] whereas complete inhibition of UPS activity was shown to occlude synaptic compensation [27]. The integral role of *de novo* translation in synaptic homeostasis was recently highlighted when proteomic analysis of neurons undergoing upscaling and downscaling revealed a remarkable diversity of newly synthesized proteins. Of particular interest was the significant enrichment in the expression of the proteasome core complex during downscaling [28,29]. The demand for the translation of proteasome complexes implies that proteasomes work alongside translation mechanisms during downscaling. Reports documenting the co-localization of ribosomes and the proteasome in neuronal dendrites [30,31] further emphasize the possibility that these two opposing machineries physically interact within the post-synaptic compartment. The remodeling of the proteome through the dynamic regulation of protein biogenesis and degradation has been termed as cellular ‘proteostasis’ [32]. However, several questions remain unexplored in the context of cellular proteostasis during homeostatic scaling, such as a) what factor establishes the link between translation and protein degradation machineries to shape the proteome during scaling? b) Which process among translation and degradation takes precedence? c) What are the signalling mechanisms that connect events of ‘sensing’ the bicuculline-mediated hyperactivity and the final down-regulation of surface AMPARs (sAMPARs)?

Here, we demonstrate a defined mechanism of synaptic scaling accomplished through an RNA-dependent coordination between translation and proteasome-mediated degradation. We observe that isolated inhibition of either translation or proteasomal activity offsets synaptic homeostasis. Restoration of homeostasis necessitates the combination of both processes. We provide empirical evidence demonstrating that the interaction between translation and protein degradation machineries is direct and RNA dependent. This coordination is achieved when the two apparatuses are tethered to actively translating transcripts linked to miRISC. Synaptic hyperactivity causes an increased abundance of Trim32 and depletion of MOV10 in polysomes; both Trim32 and MOV10 are members of the miRISC. We find that in contexts of chronic hyperactivity, mTORC1-dependent translation of the E3 ligase Trim32 promotes the poly-ubiquitination and subsequent degradation of MOV10 by proteasome. This is triggered by the mTORC1-mediated phosphorylation of its downstream effector, p70 S6 kinase (p70 S6K). We observe that MOV10 degradation leads to enhanced translation of Arc and results in the reduced distribution of sAMPARs. Loss of MOV10 alone is sufficient to decrease the synaptic strength by reducing sAMPARs and mimic events similar to hyperactivity-driven downscaling. Notably, the observed increase in Arc expression in the context of synaptic downscaling happens *via* translation and not by transcriptional mechanisms.

## Results

### Co-dependence of protein synthesis and degradation drives synaptic homeostasis

To test the existence of coordination between translation and degradation in the regulation of synaptic homeostasis, we measured miniature excitatory postsynaptic currents (mEPSCs) from cultured hippocampal neurons (DIV 18-24) after pharmacological inhibition of protein synthesis (anisomycin, 40µM) and proteasomal activity (lactacystin, 10µM) for 24 hours. Application of either lactacystin or anisomycin increased (2.43 ± 0.43 pA, p<0.02) and decreased (5.86 ± 0.13 pA, p<0.01) mEPSC amplitude respectively. Co-application of both inhibitors restored mEPSC amplitude to that of vehicle treated neurons (Figure 1A-B). The frequency of mEPSCs remained unaltered upon inhibition of translation and proteasome blockade either alone or in combination (Figure 1C), suggesting that this could be a post-synaptic phenomenon. Our data implies that interfering with either protein synthesis or degradation disturbs the balance of synaptic activity, while blocking both synthesis and degradation altogether restores it. Next, we stimulated synaptic downscaling using bicuculline (10µM, 24 hours) and observed that, like previous reports, here too, chronic application of bicuculline lead to a significant decrease in mEPSC amplitude (5.70 ± 0.08 pA, p<0.01) without any detectable change in frequency (Figure 1D-F). The extent of decrease in mEPSC amplitude within bicuculline-treated neurons recapitulated the decrease observed in neurons where translation was blocked (bicuculline treated neuron 5.70 ± 0.08 pA decrease *vs.* anisomycin treated neuron 5.86 ± 0.13 pA decrease) (Figure 1B and 1E). We measured the mEPSC amplitude and frequency from hippocampal neurons when bicuculline was co-applied with anisomycin and lactacystin. The dual application of bicuculline and anisomycin did not result in any significant change in mEPSC amplitude when compared to neurons treated with bicuculline alone (Figure 1D-E). This confirms that, rather than inducing an additive effect, chronic inhibition of protein synthesis in itself is sufficient to induce downscaling and could potentially override the effect observed due to bicuculline. Disruption of proteasome function by lactacystin during bicuculline-treatment led to a significant increase in mEPSC amplitude (9.46 ± 0.07 pA increase as compared to bicuculline treated neurons, p<0.001) without altering frequency (Figure 1E-F). The increase was effectively more than the basal activity of vehicle-treated neurons (3.76 ± 0.08 pA, p<0.01) and mimicked the increase in mEPSC amplitude brought by lactacystin alone (Fig 1B and 1E). Although the influence of lactacystin on mEPSC amplitude is opposite to that of anisomycin, their individual effects override that of bicuculline in each condition. Co-application of both inhibitors during bicuculline-induced hyperactivation produced mEPSC amplitudes comparable to vehicle treated neurons (Figure 1E). Our data indicates that the co-inhibition of translation and degradation restricts any molecular changes away from the basal level, thus, maintaining the synaptic strength at the established physiological set point.

**Figure 1:**
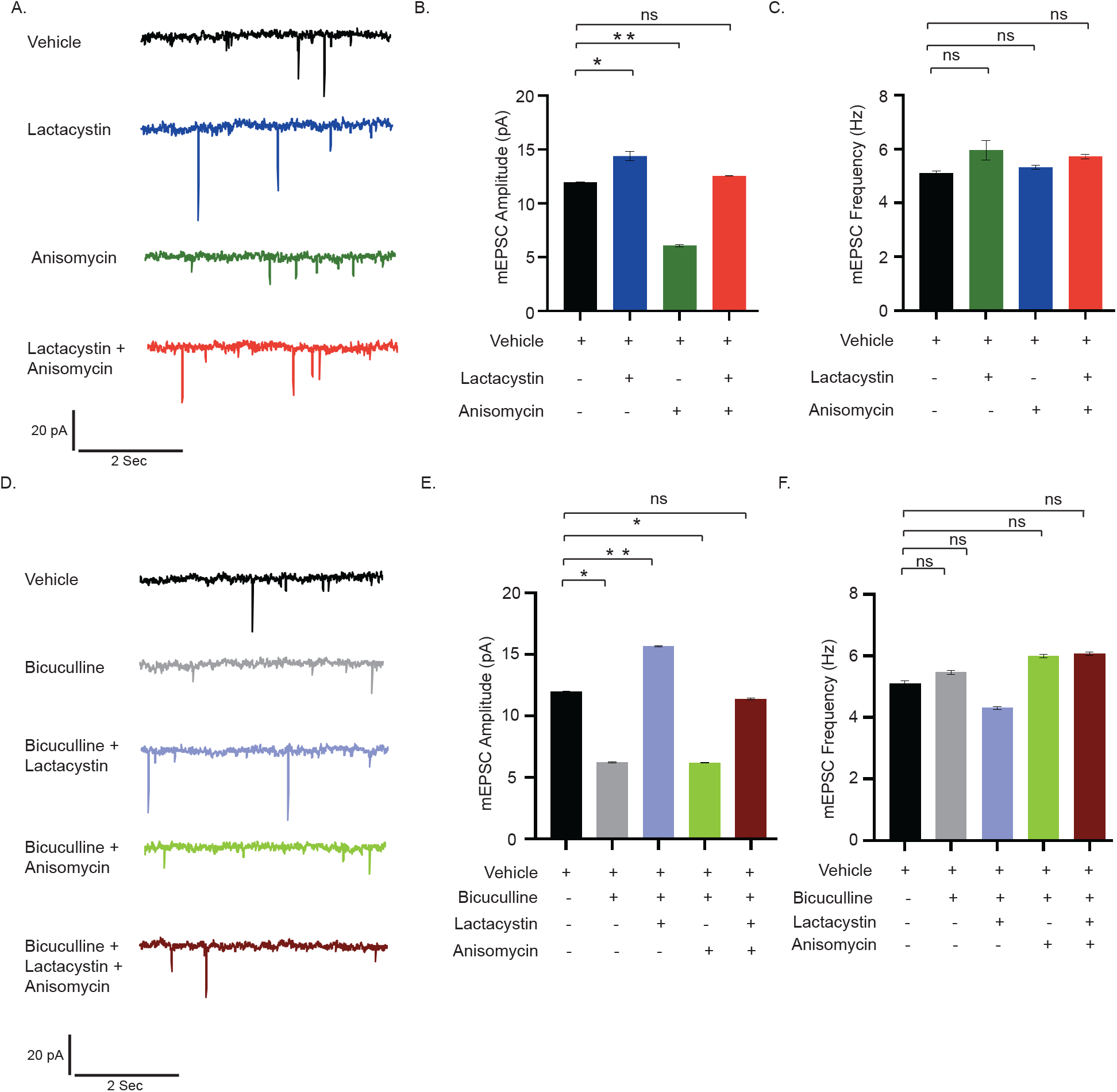
Synaptic scaling is co-­-regulated by protein synthesis and degradation. (A-C) mEPSC traces from hippocampal neurons treated with vehicle, lactacystin, anisomycin and both (A). Mean mEPSC amplitude (B) and frequency (C). n=13-15. *p<0.024, **p<0.01. ns, not significant. Data shown as Mean ± SEM. One Way ANOVA and Fisher’s LSD. Scale as indicated. (D-F) mEPSC traces from neurons treated with vehicle, bicuculline alone or in combination with lactacystin, anisomycin (D). Mean mEPSC amplitude (E) and frequency (F). n=12–16. *p<0.01, **p<0.001. ns, not significant. Data shown as Mean ± SEM. One Way ANOVA and Fisher’s LSD. Scale as indicated.

### Synchronized translation and degradation regulates AMPAR distribution during scaling

Since adjustment of synaptic strengths is directly correlated to the surface distribution of surface AMPARs (sAMPARs), we measured the surface expression of GluA1 and GluA2 (sGluA1/A2) to identify how concerted mechanisms of synthesis and degradation influence the distribution of sAMPARs during scaling. Neurons (DIV 21-24) were live-labeled using N-terminus specific antibodies against GluA1 and GluA2 following bicuculline treatment, either alone or in presence of both anisomycin and lactacystin, for 24 hours and synapses marked by PSD95. The surface expression of sGluA1/A2 in excitatory neurons was decreased following network hyperactivity (50.6 ± 6.68%, p<0.01 for sGluA1 and 26.1 ± 6.62%, p<0.01 for sGluA2) (Figure 2A-D, S1 A-B). Consistent with our electrophysiological data, inhibition of both the translation and the proteasome in bicuculline-treated neurons increased sGluA1/A2 levels (133.95 ± 8.77 %, p<0.01 for sGluA1, 53.17 ± 6.44%, p<0.001 for sGluA2) when compared to neurons treated with bicuculline alone (Figure 2C-D, S1A-B). Thus our data indicates that a dual inhibition of protein synthesis and degradation restores the synaptic sGluA1/A2 following network hyperactivity.

**Figure 2:**
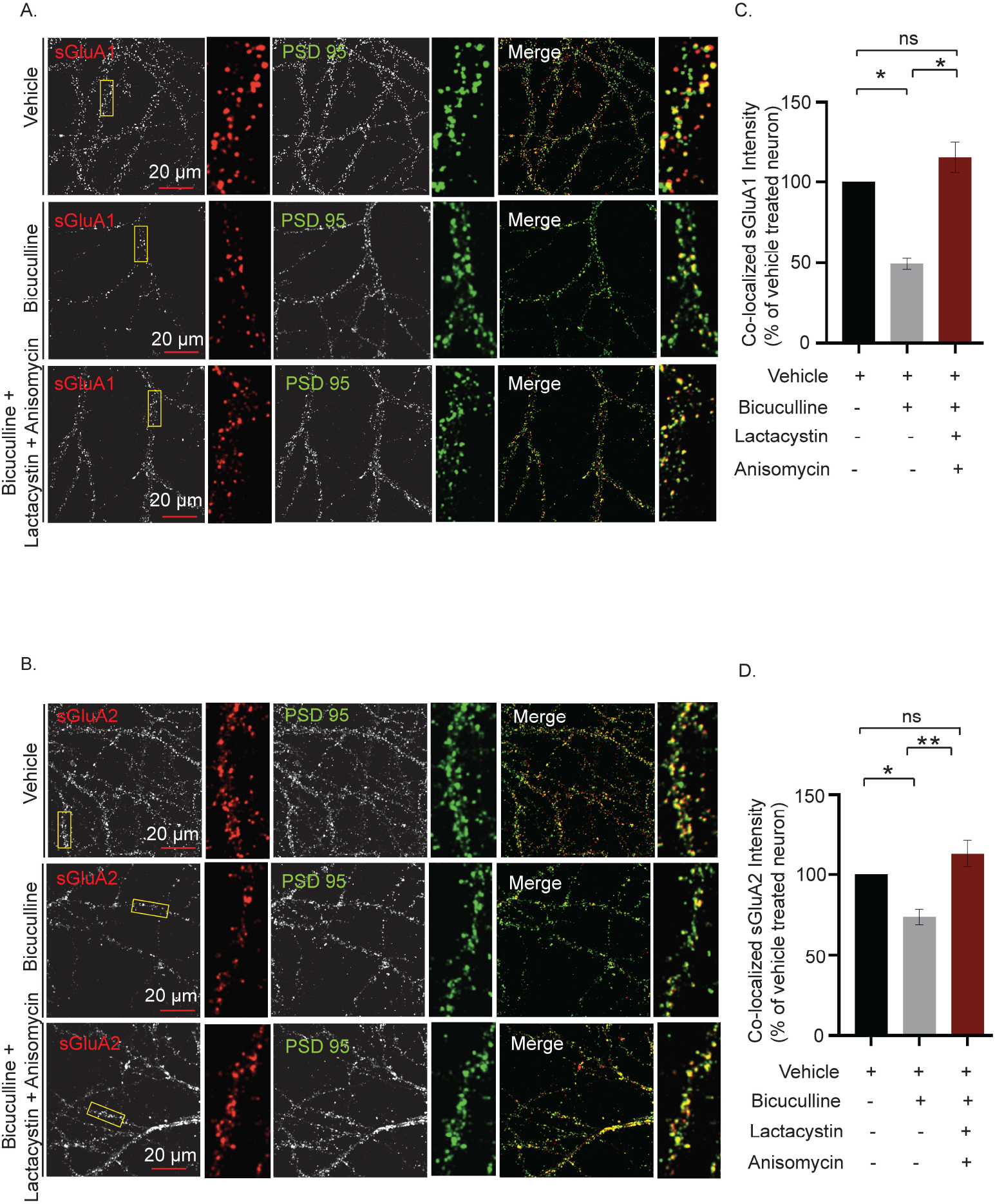
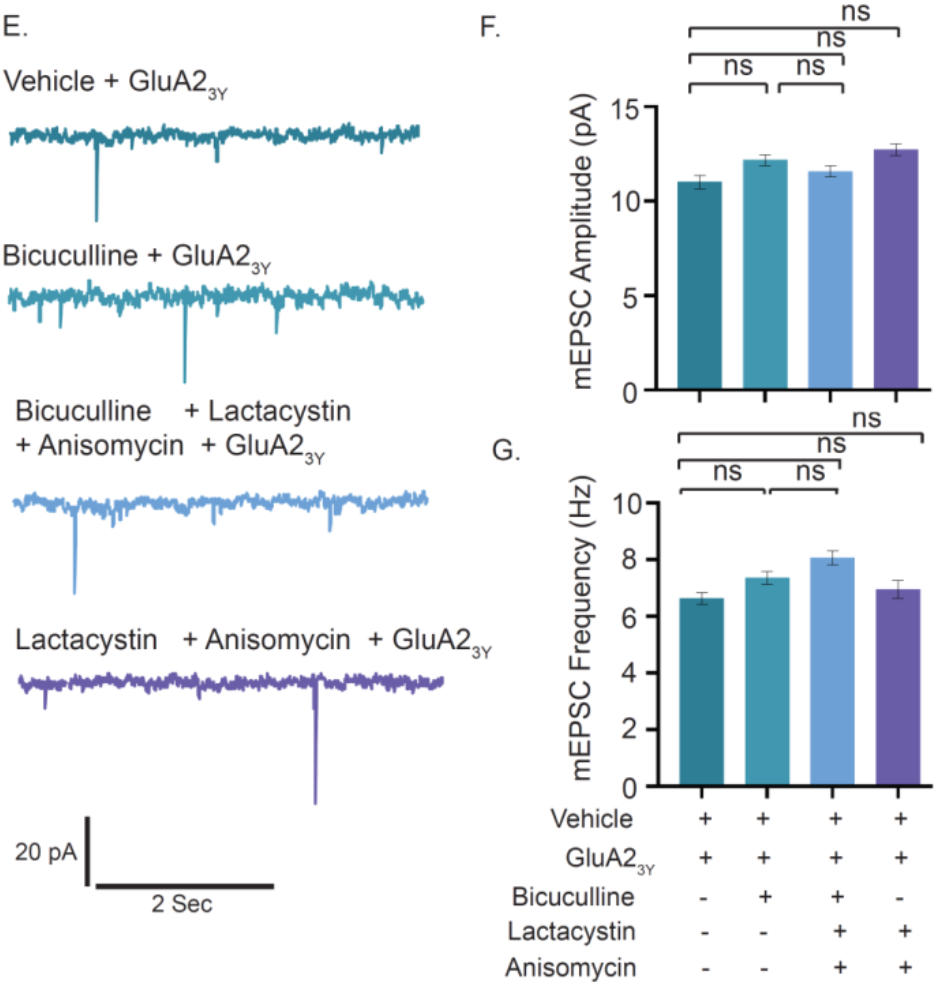
Co-­-inhibition of protein synthesis and degradation restores hyperactivity driven reduction of synaptic AMPAR expression. (A-D) High magnification images of sGluA1 or sGluA2 (red), PSD95 (green) and sGluA1/PSD95 (A) or sGluA2/PSD95 (B) (merged) images from neurons treated with vehicle, bicuculline alone or in combination with lactacystin and anisomycin. Normalized intensity of sGluA1 (C) or sGluA2 (D) co-localized with PSD95 particles. n=56-57, sGluA1 and n=31-63, sGluA2. *p<0.01, **p<0.001. One Way ANOVA and Fisher’s LSD. Dendrite marked in yellow box was digitally amplified. (E-G) mEPSCs traces from hippocampal neurons treated with GluA_23y_ either alone or in presence of bicuculline, lactacystin + anisomycin and bicuculline + lactacystin + anisomycin (E). Mean mEPSC amplitude (F) and frequency (G). n=10 - 13. ns, not significant. Data is shown as Mean ± SEM. One Way ANOVA and Fisher’s LSD. Scale as indicated. See also Figure S1.

To reaffirm whether AMPARs are indeed the end-point effectors of synaptic downscaling, we used GluA2_3Y_, a synthetic peptide derived from the GluA2 carboxy tail of AMPA receptors to block the endocytosis of the AMPARs [13], effectively ensuring that the number of AMPARs remain unchanged throughout 24 hours. Consistent with previous studies, no significant changes in mEPSC amplitude were detected upon inhibition of GluA2 endocytosis during chronic application of bicuculline (GluA2_3Y_ treated neuron 11.01 ± 0.36 pA *vs.* GluA2_3Y_ + bicuculline treated neuron 12.17 ± 0.28 pA, p<0.49) (Figure 2E-F).

Application of GluA2_3Y_ did not alter mEPSC amplitude as compared to vehicle treated neurons (GluA2_3Y_ treated neuron 11.01 ± 0.36 *vs.* vehicle treated neurons 11.94 ± 0.07 pA) (Figure S1C-D), nor any change observed between neurons treated with GluA2_3Y_ and those treated with both lactacystin and anisomycin in presence or absence of bicuculline (Figure 2F-G). mEPSC frequency remained unaltered throughout while mEPSC amplitude in each condition was similar to that of control neurons (Figure 2G, S1E). Collectively, these observations indicate that changes in the abundance of surface-AMPARs during scaling is facilitated by proteomic remodeling that exploits both translation and degradation processes.

### RNA-dependent co-sedimentation of the proteasome and translation regulators

The co-localization of polyribosomes and proteasomes to sites of synaptic activity [30,31] lead us to examine whether the components of the 26S proteasomal machinery could remain physically associated with actively translating transcripts in order to make the necessary proteomic changes. These components include proteins forming the 19S regulatory subunits and the 20S proteasome core. We analyzed polysomes from the hippocampus of 8-10 week old rats and assessed whether the sedimentation pattern of proteasomes matches those of actively translating, polyribosome-associated mRNA fractions. We observed that several components of the proteasomal machinery such as α7 subunit of the 20S proteasome; and Rpt1, Rpt3 and Rpt6 subunits of the 19S proteasome co-sedimented with translation initiation factors such as eIF4E and p70S6 kinase within actively translating polysomes (Figure 3A-B, S2A). We also detected the polysomal distribution of MOV10, a helicase and an RNA binding protein known to be poly-ubiquitinated upon synaptic activation, and Trim32, an E3 ligase, both components of the miRISC [33] (Figure 3A-B).

**Figure 3:**
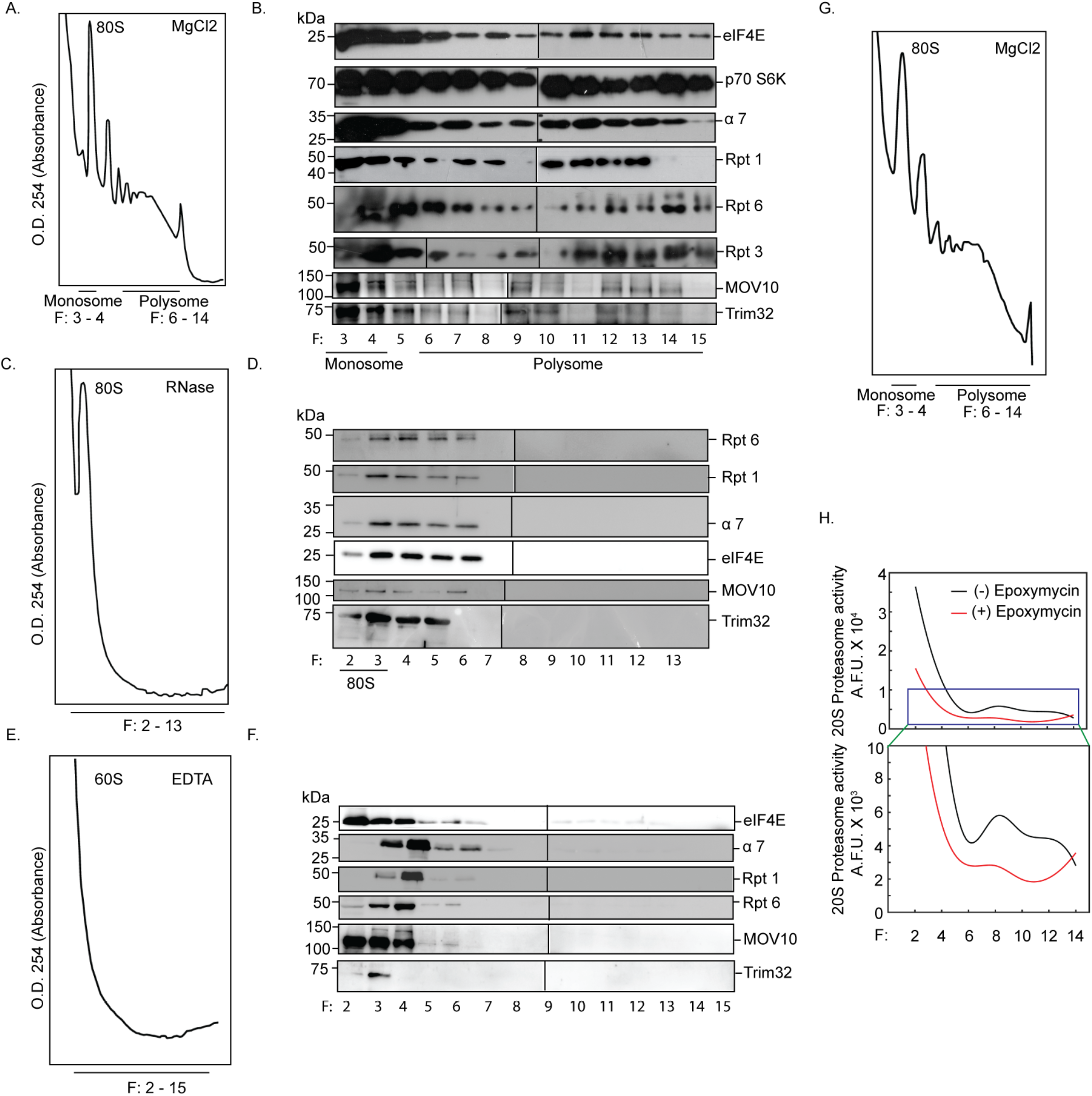
RNA-­-dependent association between active proteasomes and translating polyribosomes. (A-F) Absorbance profile at 254nm (A_254_) of fractionated cytoplasmic extracts from hippocampal tissue incubated without (A) or with RNase (C) or with EDTA (E). Monosome (80S) or 60S Ribosome or Polysome fractions as indicated. Western blot analysis of fractions without (B) or with RNase (D) or with EDTA (F) treatment showing distribution of translation regulators eIF4E and p70S6 Kinase; α7 subunit of 20S core and Rpt1, Rpt3, Rpt6 of 19S cap; miRISC proteins MOV10 and Trim32. (G-H) A_254_ profile of fractionated cytoplasmic extract (G) and quantitation of catalytic activity of proteasomes present in alternate fractions from two polysome preparations (H). See also Figure S2.

RNAse or EDTA treatment of cytoplasmic lysates prior to density gradient fractionation led to a complete collapse of the polysome profile, simultaneously shifting the sedimentation of Rpt6, Rpt1, α7, eIF4E, MOV10, Trim32 to the lighter fractions (Figure 3C-F, S2B-C). The disruption of association between the translational and proteasomal components on RNase and EDTA treatment suggests that translating transcripts are necessary to recruit the translation and proteasome machineries. These observations ruled out the possibility that the observed co-sedimentation was a result of similar densities between the protein complexes and polysomes. Trim32 and MOV10 in specific high-density sucrose fractions (fraction # 8/11/15) were not detected due to loss of proteins during the TCA precipitation step (Figure 3B). Furthermore, we saw that the polysome-associated 26S proteasome is catalytically active as detected by its ability to cleave a fluorogenic proteasome substrate that is blocked by the proteasome inhibitor epoxymycin (Figure 3G-H, S2D).

### Proteasome and the regulators of translation directly interact with each other within excitatory neurons

Whole-cell patch clamp recordings from hippocampal excitatory neurons demonstrated that the co-regulation of translation and proteasome-mediated protein degradation is necessary for synaptic homeostasis. Consistent with this observation, co-sedimentation of proteasome subunits along with polysomes linked to protein synthesis regulators and members of the miRISC led us to enquire whether components of the ternary complex directly interact with each other in excitatory neurons of the hippocampus. To evaluate this; we immunoprecipitated the 19S proteasomal complex using Rpt6 antibody from hippocampal neurons. We observed the co-precipitation of eEF2, a translation elongation factor that functions as a “sensor” of change in network activity (Figure 4A). We also found that a known regulator of the mTORC1-dependent protein synthesis; p70 S6 kinase as well as its phosphorylated form [34] co-precipitated with the 19S proteasome (Figure 4A). We further analyzed the proteins interacting with polysomes within excitatory neurons by expressing haemagglutinin (HA)-tagged ribosomal protein Rpl22 (HA-Rpl22) that gets incorporated into polysomes [35,36] (Figures 4B-D, S2E). We reasoned that the analysis of HA-Rpl22-affinity purified complexes would confirm whether the polysome-associated translation and degradation machineries directly interact with each other.

**Figure 4:**
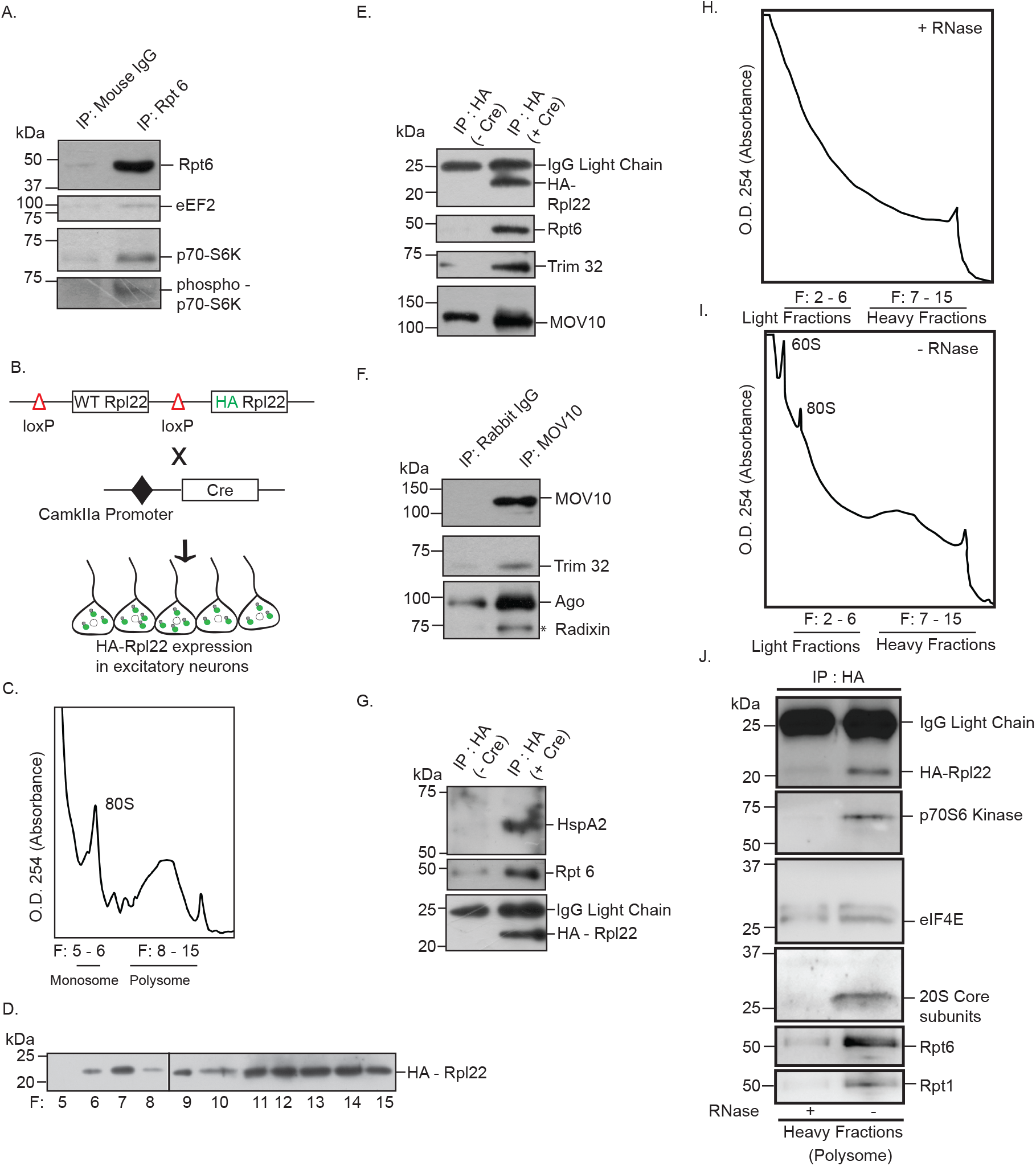
Interaction between proteasome and actively translating RNA-­-associated polyribosomes. (A) Proteasome associated protein complex was immunoprecipitated from hippocampal lysate using antibody against Rpt6 and mouse IgG. Western blot of purified protein complex performed using antibodies against eEF2, p70S6 Kinase, phospho-p70S6 kinase. (B) RiboTag mouse crossed with CamKIIa promoter-driven Cre recombinase mouse results in deletion of wild-type Rpl22 ribosomal protein and replacement of HA tagged Rpl22 in forebrain excitatory neurons. (C) A_254_ profile showing indicated fractions of Monosome and Polysome. (D) Polysome fractions showing enrichment of HA-Rpl22 as detected by western blot using antibody against HA. (E) HA-tagged Rpl22 containing polyribosome affinity purified using antibody against HA. Western blot analysis of affinity purified complex using antibodies against HA, Rpt6, Trim32 and MOV10. (F) MOV10 was immunoprecipitated from hippocampal lysate. Western blot analysis of MOV10-immunoprecipitated protein complex showed the co-precipitation of Trim32 with miRISC components MOV10 and Ago. (G) Detection of HspA2 and Rpt6 in HA affinity purified protein complex from HA-Rpl22 expressing neurons by western blot using antibody against HspA2 and Rpt6 and HA. (H – I) A_254_ profile showing indicated fractions of Monosome and Polysome obtained from cytoplasmic lysates treated with (H) or without (I) RNase prior to density gradient fractionation. (J) HA-tagged Rpl22 containing ribosomes affinity purified from heavy fractions of sucrose gradient using antibody against HA. Western blot analysis of affinity purified complex using antibodies against HA, p70 S6 kinase, eIF4E, 20S Core subunits, Rpt6 and Rpt1.

Our western blot analysis of HA-Rpl22 affinity-purified protein complex showed that Rpt6 directly interacts with Trim32 and MOV10 (Figure 4E). Immunoprecipitation of MOV10 from hippocampal neurons resulted in the co-precipitation of both Argonaute (Ago) and Trim32, confirming that the latter is an integral component of the Ago-containing miRISC (Figure 4F). We also detected the chaperone protein HspA2 in the HA-affinity purified fraction along with Rpt6 (Figure 4G), suggesting that HspA2 could tether proteasomes to actively translating transcripts. The direct interaction between components of the translation and proteasome machinery could occur without the participation of polysome-associated, translating mRNA. To evaluate whether the observed association was RNA-dependent or RNA-independent, HA-Rpl22-affinity purified protein complexes from polysome fractions of hippocampal tissue lysates treated with or without RNase were analyzed. Our western blot analysis revealed that the 20S proteasome core, Rpt6 and Rpt1 co-precipitated with eIF4E and p70 S6 kinase (Figure 4J). RNase treatment of the cytoplasmic lysate prior to density gradient fractionations prevented this interaction on the actively translating, heavier fractions of the polysome (Figure 4J). This demonstrates that polysome-associated, translating RNA act as scaffolds to facilitate the direct interaction between protein synthesis and degradation modules.

### Chronic hyperactivity in neurons regulates the distribution of factors associated with translation and the proteasomal machinery

Once we identified that members of the translation apparatus, the 26S proteasome and the miRISC remain directly associated on polysomes, we investigated the effect of prolonged neuronal hyperactivity on the polysomal distribution of these factors in the context of synaptic homeostasis. To evaluate the hyperactivity-regulated polysome association, density gradient fractions from neurons treated with either bicuculline (10µM, 24 hours) or vehicle were analyzed by western blot using antibodies against translation regulators, proteasome subunits, chaperone and members of the miRISC. We observed a relative enrichment of the translation elongation factor eEF2, mTORC1 downstream effector p70 S6 kinase as well as its phosphorylated form and the phosphorylated ribosomal protein S6 in polysome fractions following prolonged neuronal activity (Figure 5A - D). Similar to enrichment of translation regulators, we also observed an enrichment of 20S core proteasome subunits upon bicuculline treatment (Figure 5A – D). The chaperone protein HspA2 was detected in polysome fractions in both vehicle-treated and bicuculline-treated neurons (Figure 5A – D). The core component of the miRISC, Argonaute showed a relative depletion from polysome fractions in bicuculline-treated neurons as compared to vehicle-treated neurons. Furthermore, we observed that the abundance of Trim32 was enhanced (108.3 ± 7.74% increase, p<0.005) whereas MOV10 (72.37 ± 3.54% decrease, p<0.002) was depleted from polysome in response to synaptic hyperactivity (Figure 5 C –F).

**Figure 5:**
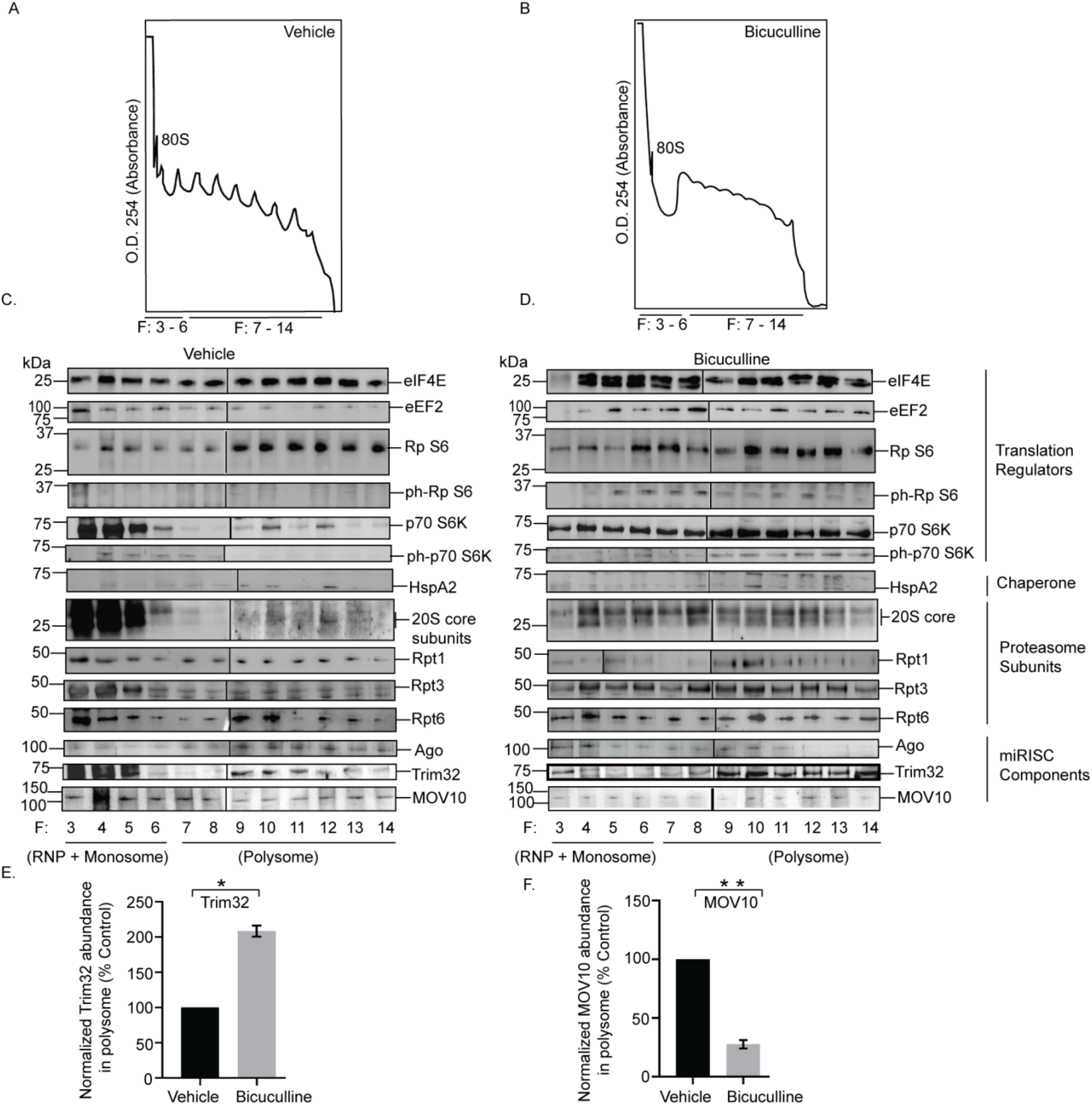
Hyperactivity-­-dependent polysome distribution of proteasome, translation regulators and component of miRISC. (A-B) Absorbance profile at 254nm (A_254_) of fractionated cytoplasmic extracts from cortical neurons treated with vehicle (A) or bicuculline (B). Monosome (80S) or Polysome fractions as indicated. Western blot analysis of fractions from vehicle (C) or bicuculline (D) treated neurons showing distribution of translation regulators eIF4E, eEF2, ribosomal protein S6 (Rp S6), phosphorylated ribosomal protein S6 (ph-Rp-S6), p70 S6 Kinase (p70 S6K), phosphorylated p70 S6 Kinase (ph-p70 S6K); chaperone protein HspA2; proteasome subunit of 20S core and Rpt1, Rpt3, Rpt6 of 19S cap; miRISC proteins Argonaute (Ago), MOV10 and Trim32. (E) Quantitation of polysome distribution of Trim32. (F) Quantitation of polysome distribution of MOV10. *p<0.005, **p<0.002. n=3, Unpaired t-test with Welch’s correction. RNP: Ribonucleoprotein. See also Figure S2.

Comprehensively, these data point toward activity-induced dynamicity in polysome association of factors regulating synaptic homeostasis.

### Translation drives proteasomal degradation to cause miRISC remodeling during synaptic downscaling

The reciprocal pattern of abundance of MOV10 and Trim32 in polysomes upon chronic hyperactivity (Figure 5E-F) and their association with Argonaute (Figure 4F), a core member of the miRISC, prompted us to analyze their expression in the context of synaptic downscaling. Bicuculline treatment of hippocampal neurons (DIV 18 - 21) enhanced (108.2 ± 7.55% increase, p<0.0001) Trim32 with a concomitant decrease (65.94 ± 2.67% decrease, p<0.001) in MOV10 (Figure 6A-C). The increase in Trim32 expression post bicuculline treatment was blocked by anisomycin and surprisingly resulted in the inhibition of MOV10 degradation (184.82 ± 10.77% protected MOV10, p<0.03) (Figure 6C). This indicates that the degradation of MOV10 is dependent on enhanced Trim32 synthesis and that Trim32 translation precedes the commencement of MOV10 degradation. Treatment with lactacystin resulted in the expected protection of MOV10 from degradation (202.41 ± 3.18% MOV10 protected, p<0.006) upon bicuculline-induced hyperactivity (Figure 6C); whereas there remained no change in the Trim32 expression levels (Figure 6B). Co-application of lactacystin and anisomycin during bicuculline-induced hyperactivity changed the expression of MOV10 and Trim32 commensurate to basal levels (Figure 6B-C). We did not observe any alteration of Argonaute expression upon prolonged bicuculline treatment of hippocampal neurons (Figure S3A-B). Chronic inhibition of protein synthesis alone, without bicuculline treatment, led to a modest but statistically significant decrease of both Trim32 (22.42 ± 0.70% decrease, p<0.001) and MOV10 (28.14 ± 0.48% decrease, p<0.0003) (Figure S3C-E), whereas chronic inhibition of the proteasome alone has no effect on Trim32 and MOV10 expression (Figure S3C-E) under basal conditions. The significant decrease of both proteins upon anisomycin treatment may be due to the combined effect of global inhibition of translation and the on-going basal level of protein degradation.

**Figure 6:**
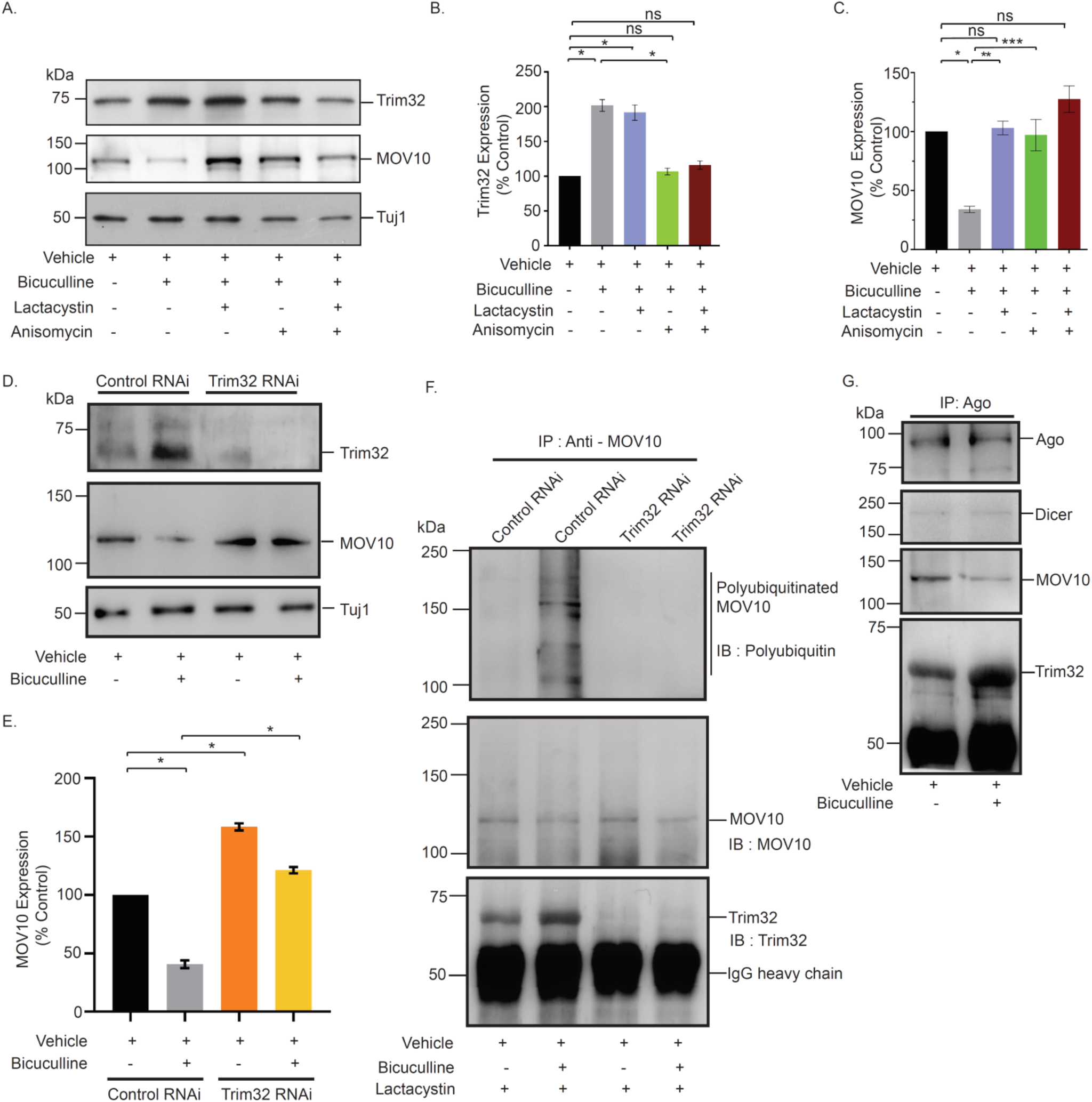
Synthesis of Trim32 facilitates polyubiquitination and subsequent degradation of MOV10 for miRISC remodeling. (A-C) Western blot analysis showing the expression of Trim32, MOV10 and Tuj1 from neurons treated with bicuculline with or without lactacystin, anisomycin or both (A). Quantitation of Trim32 (B) and MOV10 (C) expression. n=3. Data shown as Mean ± SEM. *p<0.0001 (B) and *p<0.001, **p<0.006, ***p<0.02 (C). ns, not significant. One Way ANOVA and Fisher’s LSD. (D-E) Western blot analysis of bicuculline or vehicle treated neurons infected with lentivirus expressing Trim32 or non-targetting control shRNA showing expression of Trim32, MOV10 and Tuj1 (D). Quantitation of Trim32 (n=5) (E). Data shown as Mean ± SEM. *p<0.0001. Unpaired t-test with Welch’s correction. (F) MOV10 was immunoprecipitated from hippocampal neurons treated with bicuculline or vehicle in presence of lactacystin. Polyubiquitination of MOV10 was detected by western blot analysis using an antibody against polyubiquitin conjugates (FK1). Western blot analysis of MOV10-immunoprecipitated protein complex showed the co-precipitation of Trim32 in presence or absence of bicuculline. (G) Argonaute (Ago) was immunoprecipitated from hippocampal neurons treated with bicuculline or vehicle. Western blot analysis of immunoprecipitated Argonaute complex showed the co-precipitation of miRISC proteins Dicer, MOV10 and Trim32.

The observed reciprocity between MOV10 and Trim32 expression levels on chronic bicuculline treatment led us to analyze whether Trim32 is the sole E3-ligase responsible for the UPS-mediated degradation of MOV10. Knockdown of Trim32 prevented the bicuculline-induced degradation of MOV10 (199.41 ± 0.69% protected, p<0.0001) (Figure 6D-E). Moreover, loss of Trim32 alone enhanced the expression of MOV10 (58.3 ± 3.09% increase, p<0.0001) as compared to basal level (Figure 6D-E). We immunoprecipitated MOV10 from bicuculline or vehicle treated hippocampal neurons expressing shRNA against Trim32 or control shRNA. MOV10 ubiquitination was analyzed by western blot using an antibody that specifically recognizes polyubiquitin conjugates. We observed that bicuculline-induced polyubiquitination of MOV10 was abrogated by Trim32 knockdown (Figure 6F). These observations indicate that a) the translation of Trim32 is a prerequisite for the degradation of MOV10 by proteasome during synaptic downscaling and b) Trim32 is the only E3 ligase necessary and sufficient for MOV10 ubiquitination.

Bicuculline-induced changes in Trim32 and MOV10 levels without any change in Argonaute suggest that during synaptic hyperactivity, miRISC remodeling could occur due to Trim32-translation-dependent MOV10 degradation. To confirm this, we have immunoprecipitated Argonaute and analyzed the association of key components of the miRISC post bicuculline treatment for 24 hours. We found a relative depletion of MOV10 and enrichment of Trim32 within the silencing complex upon synaptic hyperactivity (Figure 6G). However, the association of Dicer, another member of the miRISC, remains unaffected (Figure 6G). Comprehensively, our data point towards chronic network hyperactivity-induced miRISC remodeling that alters the expression of miRISC members, Trim32 and MOV10.

### Translation of Trim32 during chronic hyperactivity is mTORC1 dependent

Co-precipitation of the downstream effectors of the mTORC1 signaling cascade with the 26S proteosomal subunit Rpt6 (Fig 4A) and the activity-dependent differential distribution of these effectors in polysome (Fig 5C-D) led us to examine whether mTORC1 signaling plays a role in causing synaptic downscaling in response to chronic hyperactivity. Bicuculline-treatment of hippocampal neurons in the presence of rapamycin (100nM, 24 hours), a selective inhibitor of mTORC1, completely abolished the chronic hyperactivity-driven Trim32 synthesis (16.48 ± 10.33% increase as compared to control, p=0.45) and consecutive MOV10 degradation (8.19 ± 2.81% decrease as compared to control, p=0.06) (Figure 7A-C). Rapamycin treatment alone did not alter the expression patterns of Trim32 and MOV10 (Figure 7A-C). This led us to hypothesize that mTORC1 pathway acts upstream of Trim32, serving to regulate its synthesis in response to chronic bicuculline treatment. Consistent with our biochemical data, we observed that co-incubation of rapamycin and bicuculline prevented the decrease in mEPSC amplitude (2.76 ± 0.13 pA increase as compared to bicuculline treated neurons, p<0.01) but not frequency (Figure 7D-F). Just as above, rapamycin treatment alone has no effect, indicating that chronic hyperactivity acts as a triggering point for mTORC1 activation (Figure 7D-F) and this subsequently plays a role in driving Trim32 translation.

**Figure 7:**
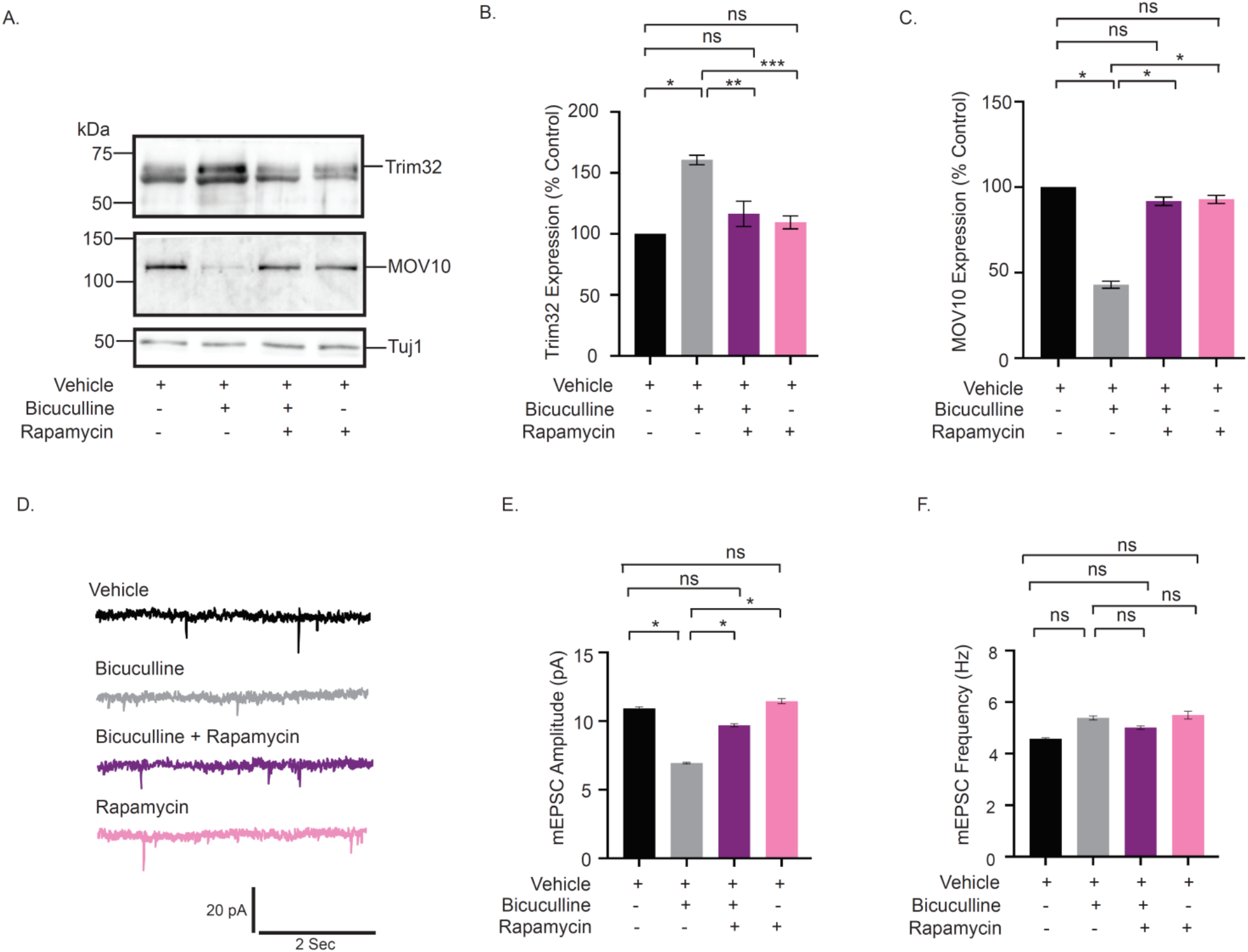
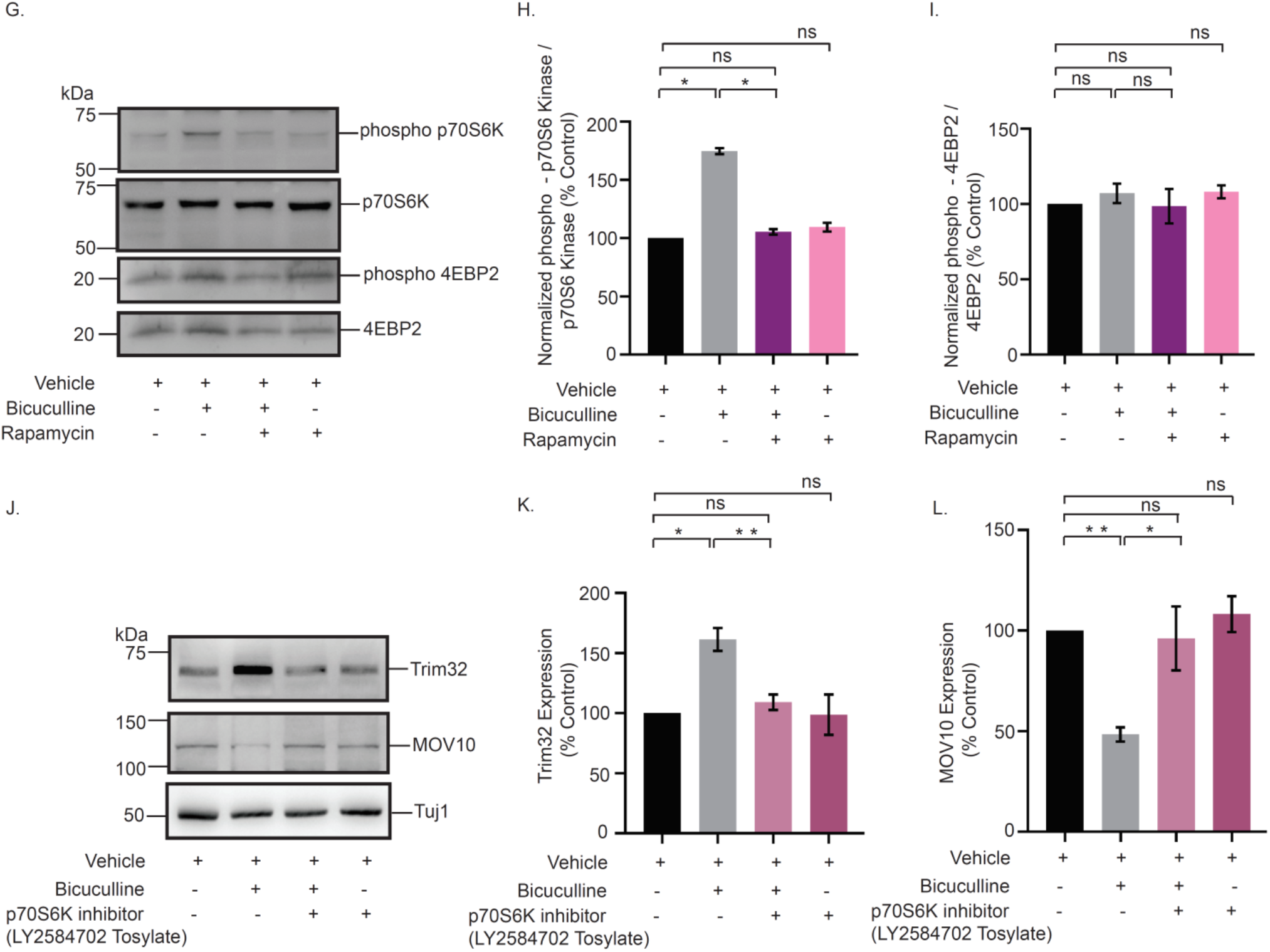
mTORC1 regulates Trim32 synthesis and consequent MOV10 degradation during synaptic downscaling. (A-C) Western blot analysis from neurons treated with bicuculline, rapamycin or both showing expression levels of Trim32, MOV10 and Tuj1 (A). Quantitation of Trim32 (B) MOV10 (C) expression. Data shown as Mean ± SEM. n=5. *p<0.0001, **p<0.0007, ***p<0.0001 (B) and *p<0.001 (C). ns, not significant. One Way ANOVA and Bonferroni’s correction. (D-F) mEPSC traces from neurons treated with vehicle, bicuculline, rapamycin or both (D). Mean mEPSC amplitude (E) and frequency (F). n=8-9. *p<0.01. ns, not significant. Data shown as Mean ± SEM. One Way ANOVA and Fisher’s LSD. (G-I) Western blot analysis from neurons treated with bicuculline, rapamycin or both showing phosphorylation and total expression of p70 S6 Kinase and 4EBP2 (G). Quantitation of p70 S6 Kinase phosphorylation (H) 4EBP2 phosphorylation (I). n=4. *p<0.0001. ns, not significant. Data shown as Mean ± SEM. One Way ANOVA and Fisher’s LSD. (J-L) Western blot analysis from neurons treated with bicuculline, p70 S6 kinase inhibitor LY2584702 Tosylate or both showing expression of MOV10 and Trim32 (J). Quantitation of Trim32 (K) and MOV10 (L). n=5. *p<0.003, **p<0.0002 (K) **p<0.0001, *p<0.02 (L). ns, not significant. Data shown as Mean ± SEM. One Way ANOVA and Fisher’s LSD.

mTORC1-mediated translation is regulated by the phosphorylation of two divergent downstream effectors, eukaryotic translation initiation factor 4E-binding protein 2 (4EBP-2) and p70 S6 Kinase (p70 S6K) [37]. We examined the phosphorylation status of these two downstream effectors following bicuculline-induced hyperactivity. Chronic bicuculline treatment leads to a significant enhancement of p70 S6 kinase phosphorylation (74.66 ± 2.61 % increase, p<0.0001) which was blocked by rapamycin (Figure 7G-H). However, bicuculline treatment did not affect the phosphorylation of 4EBP2 (Figure 7G and 7I).

We used a specific inhibitor of p70 S6K, LY2584702 Tosylate (2 µM, 24 hours), in primary hippocampal neurons to evaluate the role of p70 S6K activity on Trim32 and MOV10 expression in the context of synaptic downscaling. Similar to rapamycin treatment, the inhibition of p70 S6K phosphorylation prevented Trim32 translation and the subsequent degradation of MOV10 upon bicuculline-induced hyperactivity (Figure 7J-L). These observations showed that the activation of p70 S6K by mTORC1 is a key determinant regulating the concerted translation of Trim32 and the degradation of MOV10 in downscaling.

### MOV10 degradation is sufficient to invoke downscaling of AMPARs

MOV10 degradation in response to chronic bicuculline treatment made us enquire whether its loss alone was sufficient to cause pervasive changes in the miRISC to effectuate synaptic downscaling. We used lentivirus-mediated RNAi of MOV10 to mimic hyperactivity-driven MOV10 degradation. Intensity of sGluA1/A2 puncta that co-localized with PSD95 was analyzed following MOV10 knockdown (DIV21-24). We observed that loss of MOV10 reduced the expression of sGluA1 (35.03 ± 9.35 % for shRNA#1, p<0.01 and 58.38 ± 10.27 % for shRNA#2, p<0.01) and sGluA2 (49.4 ± 12.9% for shRNA#1, p<0.01) at the synapses (Figure 8A-D, S4), that recapitulated the re-distribution of sGluA1/sGluA2 in neurons under chronic bicuculline treatment (Figure 2C-D). The knockdown of MOV10 reduced mEPSC amplitude (3.43± 0.16 pA for shRNA#1 and 4.35± 0.14 pA for shRNA#2, p<0.01) but not frequency (Figure 8E-G), an observation that mirrors synaptic downscaling (Figure 1E).

**Figure 8:**
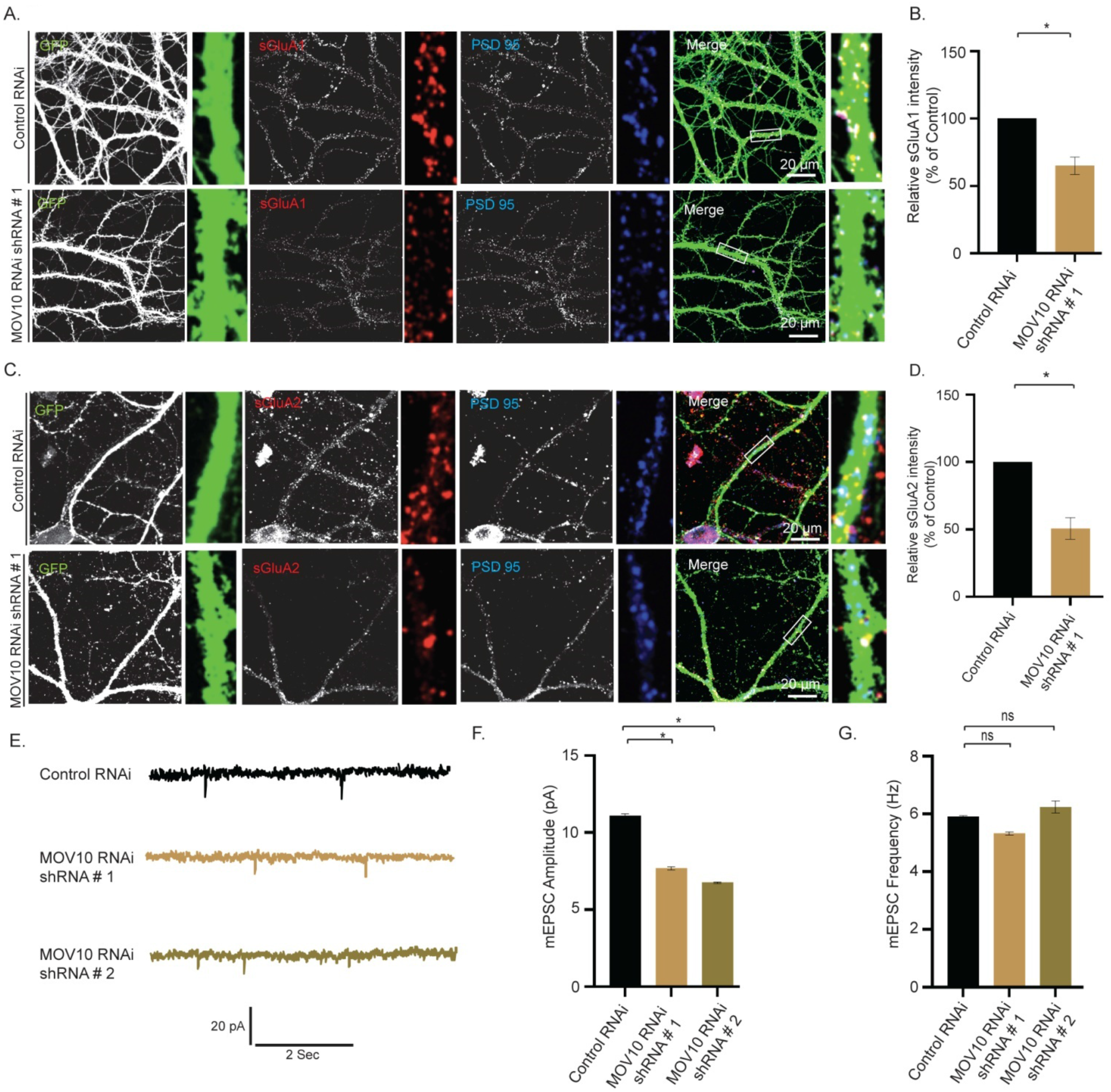
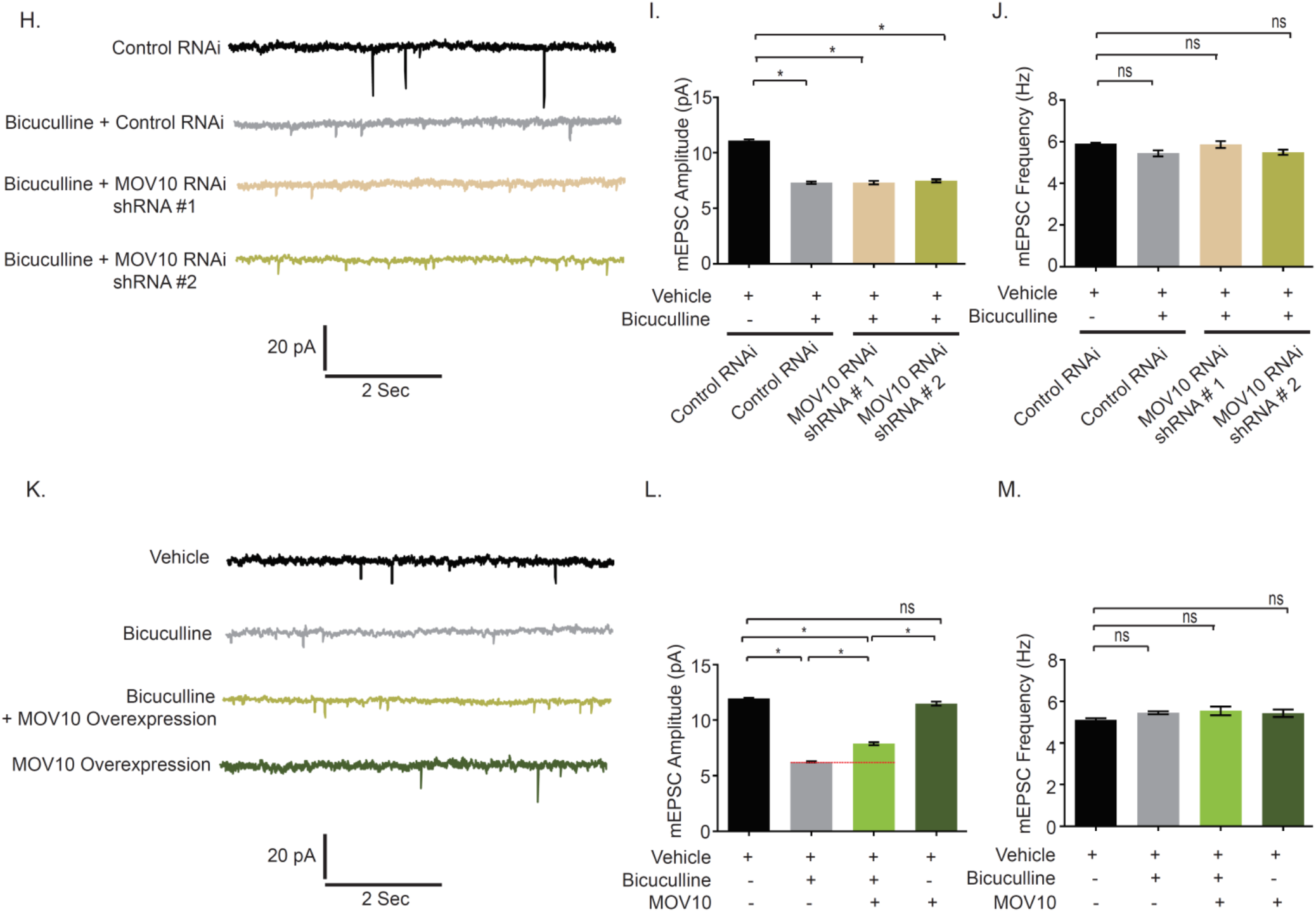
MOV10 modulates abundance of sAMPARs during synaptic downscaling. (A-D) High magnification images of neurons transduced with lentivirus co-expressing EGFP and shRNA against MOV10 or non-targeting shRNA showing expression of sGluA1 (A) or sGluA2 (C) (red), PSD95 (blue), GFP (green) and GFP/sGluA1/PSD95 or GFP/sGluA2/PSD95 (merged). Quantitation of normalized intensity of synaptic sGluA1 (B) or sGluA2 (D). n=26 - 30, GluA1; n=12 – 15, GluA2. Data shown as Mean ± SEM. *p<0.01. One Way ANOVA and Fisher’s LSD. Dendrite marked in white box was digitally amplified. See also Figure S4. (E-G) mEPSC traces from neurons transduced with shRNAs against MOV10 or control shRNA (E). Mean mEPSC amplitude (F) and frequency (G). n=12 – 13. *p<0.01. ns, not significant. Data shown as Mean ± SEM. One Way ANOVA and Fisher’s LSD. (H-J) mEPSC traces from bicuculline or vehicle treated neurons transduced with shRNAs against MOV10 or control shRNA (H). Mean mEPSC amplitude (I) and frequency (J). n=12 – 18. *p<0.0001. ns, not significant. Data shown as Mean ± SEM. One Way ANOVA and Fisher’s LSD. (K-M) mEPSC traces from bicuculline treated neurons overexpressing myc-tagged MOV10 (K). Mean mEPSC amplitude (L) and frequency (M). n=13 – 15. *p<0.0001. ns, not significant. Data shown as Mean ± SEM. One Way ANOVA and Fisher’s LSD. Dotted line indicates the difference in mEPSC amplitude recorded from neurons overexpressing MOV10 or control neurons following bicuculline treatment. See also Figure S6.

We examined whether MOV10 knockdown could further influence synaptic strength effectuated by bicuculline. Bicuculline treatment did not further decrease the amplitude of mEPSCs recorded from neurons expressing MOV10 shRNAs (Figure 8H – I). This observation indicates that effect of MOV10 knockdown overrides the effect of bicuculline. We then overexpressed myc-tagged MOV10 in hippocampal neurons and measured their synaptic activity. Bicuculline treatment led to an enhancement of mEPSC amplitude (1.64 ± 0.14 pA increase, p<0.0001) (Figure 8K – L) in neurons overexpressing MOV10 but not mEPSC frequency (Figure 8K and 8M). Our data shows that the overexpression of MOV10 partially occludes synaptic downscaling.

### Trim32 is required for AMPAR-mediated downscaling

We have examined the role of Trim32 in bicuculline-induced downscaling by measuring synaptic activity following the loss of Trim32. Trim32 knockdown in bicuculline-treated neurons led to an increase in mEPSC amplitude (2.82 ± 0.23 pA increase, p<0.0001), but did not show any change in frequency as compared to neurons incubated with bicuculline alone (Figure 9A-C). A modest but significant increase in mEPSC amplitude (1.65± 0.25 pA increase, p<0.0001) was also detected from neurons expressing Trim32 shRNA as compared to control shRNA without any activity (Figure 9A-B). We presume that this could be due to an increase in basal MOV10 expression following Trim32 knockdown (Figure 6D-E). Consistent with patch clamp recording data from MOV10 overexpressing neurons, a partial occlusion of bicuculline-induced downscaling by Trim32 knockdown establishes the requirement of Trim32-MOV10 axis in synaptic scaling. We examined the involvement of sAMPARs in regulation of Trim32-mediated downscaling. We observed that Trim32 knockdown in bicuculline-treated neurons enhanced both sGluA1 (31.35 ± 6.65% increase, p<0.04) (Figure 9D – E, 9H) and sGluA2 (88.99 ± 5.52% increase, p<0.0005) (Figure 9F – G, 9I) as compared to neurons treated with bicuculline alone.

**Figure 9:**
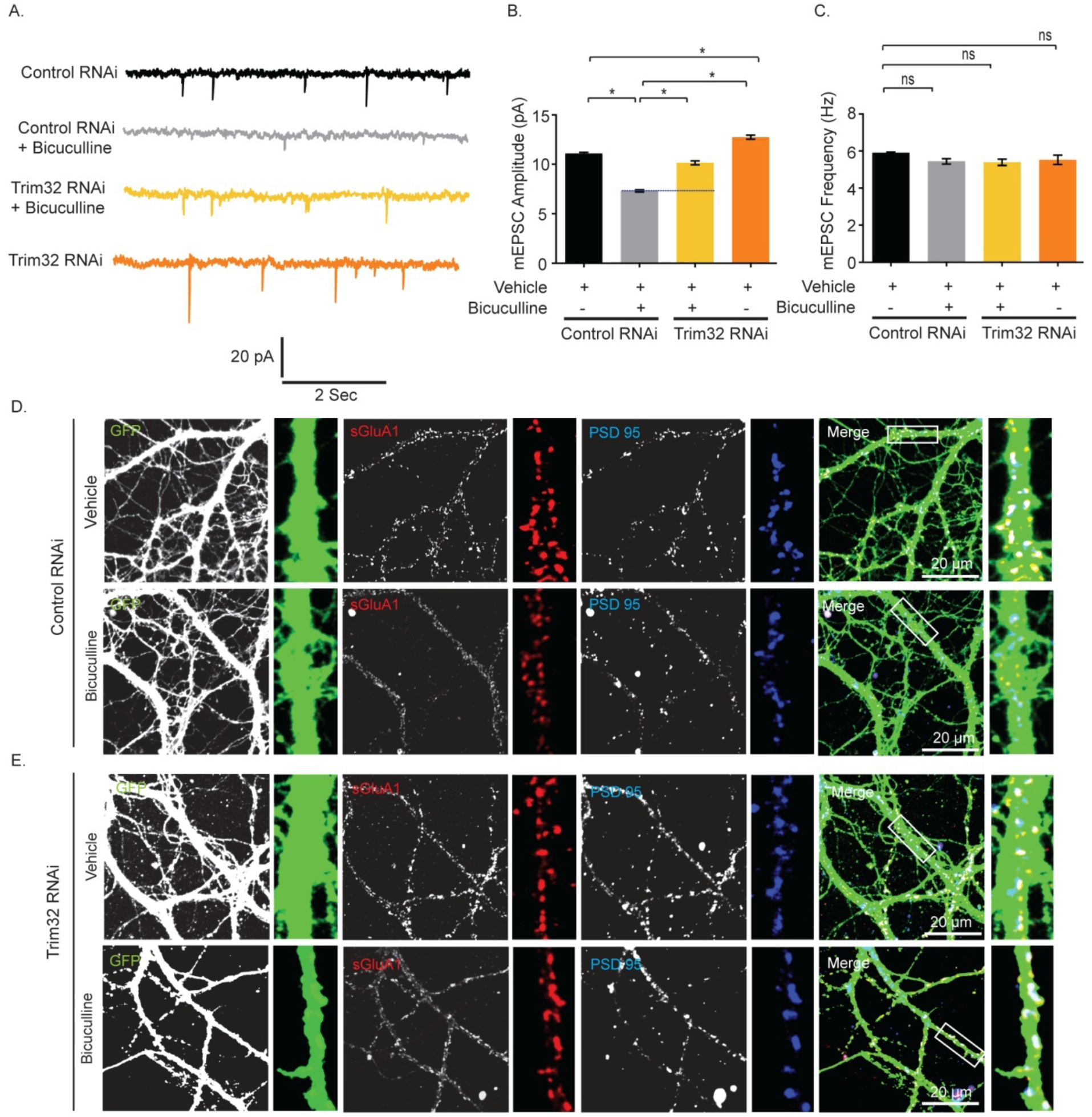
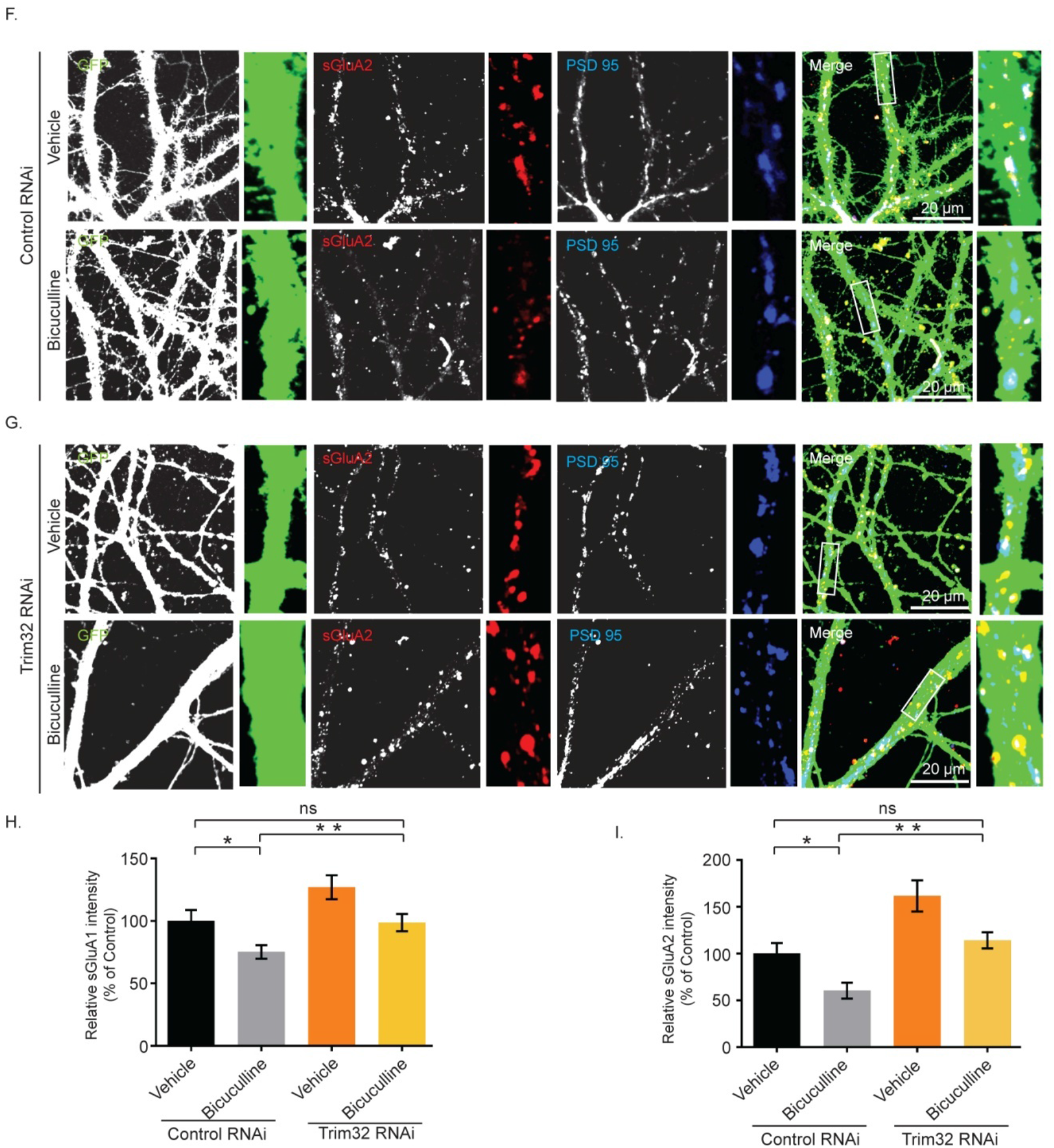
Trim32 -mediated synaptic downscaling occurs *via* modulation of sAMPARs. (A-C) mEPSC traces from bicuculline or vehicle treated neurons transduced with shRNAs against Trim32 or control shRNA (A). Mean mEPSC amplitude (B) and frequency (C). n=12 – 16. *p<0.0001. ns, not significant. Data shown as Mean ± SEM. One Way ANOVA and Fisher’s LSD. Dotted line indicates the difference in mEPSC amplitude recorded from bicuculline treated neurons transduced with Trim32 shRNA or control shRNA. (D-I) High magnification images of bicuculline or vehicle treated neurons transduced with lentivirus co-expressing EGFP and shRNA against Trim32 or non-targetting shRNA showing expression of sGluA1 (D-E) or sGluA2 (F-G) (red), PSD95 (blue), GFP (green) and GFP/sGluA1/PSD95 or GFP/sGluA2/PSD95 (merged). Quantitation of normalized intensity of synaptic sGluA1 (H) or sGluA2 (I). n=30 - 40 for GluA1; n=23 – 36 for GluA2. Data shown as Mean ± SEM. *p<0.02, **p<0.0001 for sGluA1. *p<0.006, **p<0.0005 for sGluA2. One Way ANOVA and Fisher’s LSD. Dendrite marked in white box was digitally amplified. See also Figure S5.

Similar to increase in mEPSC amplitude following Trim32 knockdown, our data showed an increase in sGluA1 (27.009 ± 7.72%, p<0.04) and sGluA2 (61.63 ± 10.47%, p<0.007) in Trim32 RNAi neurons. (Figure 9H – I). Comprehensively, our data demonstrates a regulatory control of synaptic downscaling by the Trim32-MOV10 axis that modulates sAMPAR expression.

### Trim32-MOV10 axis regulates Arc expression during synaptic downscaling

How MOV10 degradation leads to the removal of sAMPARs to regulate synaptic downscaling? Arc/Arg3.1, an immediate early gene, has been shown to be dynamically regulated by chronic changes in synaptic activity, and evokes synaptic scaling [38]. Overexpression of Arc decreases sAMPARs *via* endocytosis whereas its knockdown increases them [39]. Arc expression has been shown to be regulated by diverse mechanisms including translational control that involves miRNAs [40,41,42]. Since we found a decrease of sAMPAR distribution on MOV10 knockdown, we investigated the correlation between MOV10 and Arc expression. We observed that the loss of MOV10 enhanced Arc (113.1 ± 15.7% increase, p<0.002 for shRNA # 1 and 173.8 ± 7.45% increase, p<0.0001 for shRNA # 2) (Figure 10A-B). The extent of increase in Arc protein was commensurate with the degree of MOV10 knockdown (Figure 10A). We also observed that this differential enhancement of Arc protein was reflected in the proportionate removal of sAMPARs and the concomitant decrease in mEPSC amplitude (Figure 8F). Bicuculline-induced chronic hyperactivity, which degrades MOV10, also enhanced Arc expression (132.1 ± 27.45% increase, p<0.04) (Figure 10C-D). This activity-driven increase was blocked by the inhibition of mTORC1 with rapamycin (100nM, 24 hours) whereas rapamycin treatment alone had no effect (Figure 10C-D).

**Figure 10:**
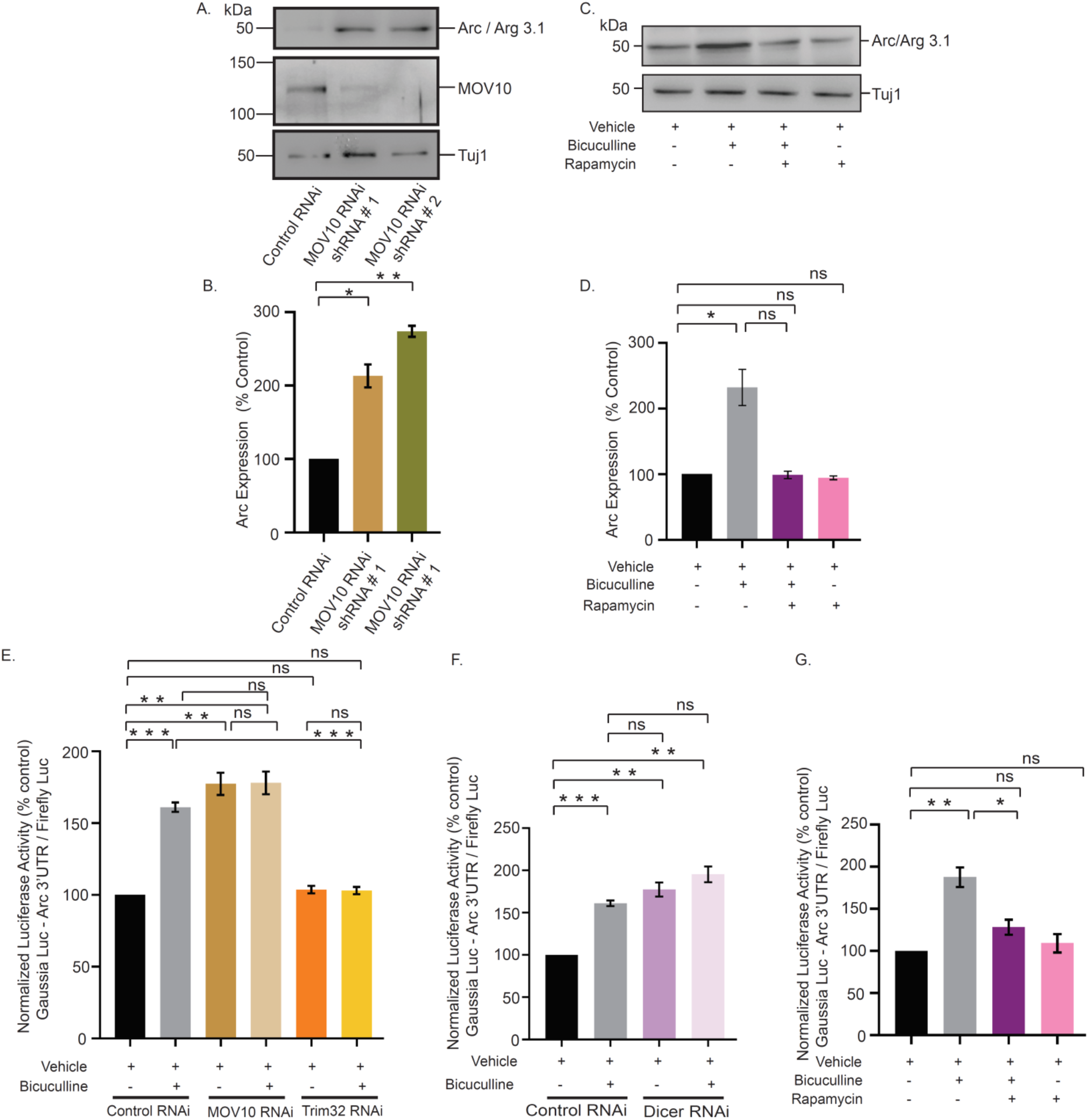
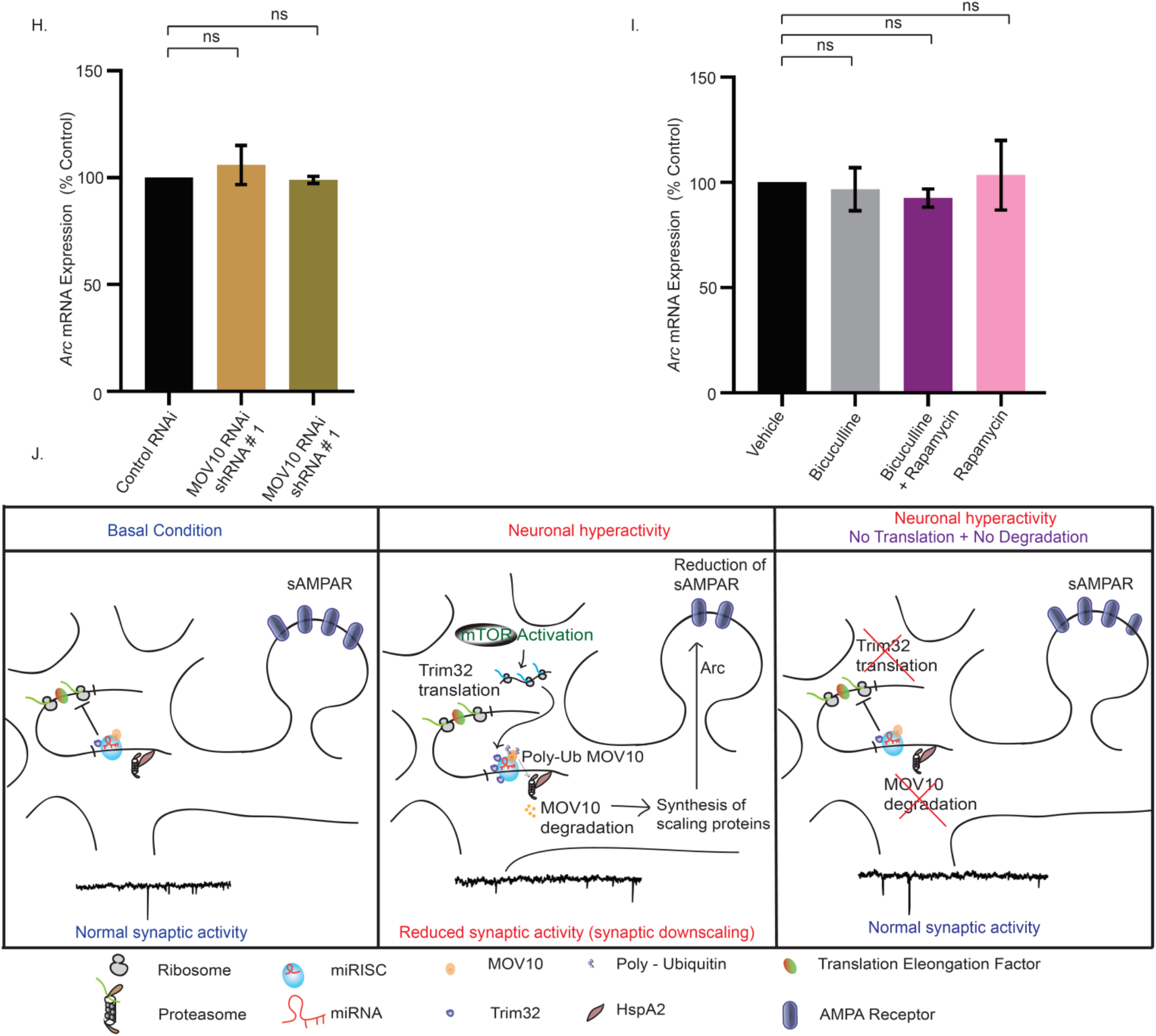
mTORC1-­-mediated regulation of Arc expression upon chronic hyperactivity involves MOV10. (A-B) Western blot analysis showing the Arc protein level after MOV10 knockdown in neurons infected with lentivirus expressing two different shRNAs against MOV10 (A). Quantitation of Arc expression (B). n=6.* p<0.002, **p<0.0001. Data shown as Mean ± SEM. One Way ANOVA and Fisher’s LSD. (C-D) Western blot analysis of neurons treated with bicuculline in presence or absence of rapamycin showing the expression of Arc protein (C). Quantitation of Arc expression (D). n=3. *p<0.04. Data shown as Mean ± SEM. One Way ANOVA and Fisher’s LSD. (E-F) Quantitation of luciferase reporter expression from vehicle or bicuculline-treated neurons transduced with lentivirus expressing shRNA against MOV10 and Trim32 (E), Dicer (F). n=4. **p<0.002, ***p<0.0003, ns, not significant. Data shown as Mean ± SEM. One Way ANOVA and Fisher’s LSD. See also Figure S6. (G) Quantitation of luciferase reporter expression from vehicle or bicuculline-treated neurons in presence or absence of rapamycin. n=7. **p<0.00, *p<0.04, ns, not significant. Data shown as Mean ± SEM. One Way ANOVA and Fisher’s LSD. (H-I) Quantitation of qPCR analysis of Arc mRNA expression in neurons expressing shRNA against MOV10 or control shRNA. n=3. ns, not significant. Data shown as Mean ± SEM. One Way ANOVA (H). Quantitation of qPCR analysis of Arc mRNA expression in bicuculline-treated neurons in presence or absence of rapamycin (I). n=4. ns, not significant. Data shown as Mean ± SEM. One Way ANOVA (I). (J) Schematic representation showing maintenance of homeostatic synaptic activity by coordinated control of protein synthesis and degradation that modulates composition of miRISC.

Arc expression has been shown to be regulated by both transcriptional and post-transcriptional control [42]. Post-transcriptional control of Arc expression is regulated by its 3’UTR that contains multiple miRNA binding sites [40]. We explored the mechanism of mTORC1-dependent Arc expression involving Trim32-MOV10 axis. We used an Arc-3’UTR fused luciferase reporter (Arc-Luc) to assess Arc expression from bicuculline-treated neurons following Trim32 and MOV10 knockdown. We found that the bicuculline-induced enhancement of reporter activity (61.1 ± 3.24% increase, p<0.0003) was prevented by Trim32 knockdown (Figure 10E) whereas MOV10 knockdown alone was sufficient to increase reporter activity under basal conditions (77.34 ± 7.78% increase, p<0.002). Loss of MOV10 in bicuculline-treated neurons did not enhance Arc-Luc reporter activity further as compared to bicuculline-treated neurons (Figure 10E). This observation is consistent with our electrophysiology data demonstrating that MOV10 knockdown in bicuculline-treated neurons did not elicit further reduction in mEPSC amplitude when compared to neurons treated with bicuculline alone (Figure 8I).

Since Dicer is a key component of the miRISC, its knockdown would definitely result in pervasive miRISC remodeling and allow us to confirm whether miRISC remodeling has a role in regulating Arc expression. Hence, we tested Arc-Luc reporter activity following Dicer knockdown and found that similar to loss of MOV10, Dicer knockdown alone was sufficient to enhance Arc-Luc reporter activity under basal conditions (77.4 ±8.31% increase, p<0.002) (Figure 10F) and this enhancement was comparable to bicuculline-induced reporter activity (77.4 ± 8.31 % increase in Dicer knockdown neurons *vs.* 61.1 ± 3.24% increase in bicuculline-treated neurons) (Figure 10F). Similar to our western blot data (Figure 10C-D), we observed that the bicuculline-induced enhancement of Arc-Luc activity (87.6 ± 11.64% increase, p<0.002) was prevented by the application of rapamycin. Application of rapamycin alone did not show any change. (Figure 10G).

To explore whether Arc was regulated by transcription during synaptic downscaling, we checked Arc mRNA expression in bicuculline treated neurons after MOV10 knockdown and after mTORC1 inhibition. qRT-PCR analysis showed no detectable change in Arc transcript levels upon MOV10 knockdown or bicuculline-induced hyperactivity in presence or absence of rapamycin (Figure 10I). Taken together, our data demonstrates that the bicuculline-induced downscaling of synaptic strength occurs *via* an mTORC1-mediated Trim32 translation-dependent MOV10 degradation involving removal of sAMPARs *via* Arc (Figure 10J).

## Discussion

Here we provide empirical evidence emphasizing that synchrony between protein synthesis and proteasomal activity is critical to establish homeostasis at synapses. We used a paradigm of chronic network hyperactivity to invoke downscaling and determined that a) translation and degradation apparatuses directly interact with each other and are tethered together by RNA scaffolds; b) it is the translation of Trim32 that drives the degradation of MOV10 to cause miRISC remodeling, thus the current paradigm is an example of translation preceding degradation; c) miRISC is a key node in the translation-degradation axis, with the mTORC1-p70 S6K pathway being the upstream signaling component and a part of the ‘sensor’ machinery, and Arc-induced removal of sAMPARs being the final effectors of downscaling.

### Co-regulation of protein synthesis and degradation drives AMPAR-mediated synaptic downscaling

We find that chronic perturbation of either translation or proteasomal activity occludes synaptic homeostasis, whereas homeostasis remains unperturbed when there is simultaneous inhibition of both (Figure 1). Chronic application of bicuculline along with either lactacystin or anisomycin leads to alterations of mEPSC amplitude that exactly mirror observations where bicuculline is absent (Figure 1B *vs* Figure 1E). Thus, the effects of bicuculline-induced changes to the existing proteome are overshadowed by those accomplished by the individual action of the proteasome or the translation machinery (Figures 1). The importance of these observations is multi-faceted; it establishes that, i) congruent protein synthesis and degradation pathways regulate synaptic scaling; ii) the constancy of the proteomic pool in the presence of lactacystin and anisomycin renders the effect of any network destabilizing stimuli like bicuculline to be redundant, and iii) bicuculline induced changes in the proteome predominantly affects the physiology of the post-synaptic compartment.

Our observations echo previous findings in Hebbian plasticity; wherein, protein synthesis during LTP/LTD was required to counter the changes in the proteomic pool triggered by protein degradation. The blockade of L-LTP accomplished by inhibiting protein synthesis was revoked on the simultaneous application of proteasomal blockers and translational inhibitors [43]. Abrogation of proteasomal activity allowed mGluR-dependent LTD to proceed when protein synthesis was co-inhibited [44]. These observations emphasize the existence of a proteostasis network that enable compositional changes to the proteome in contexts of acute or chronic changes in synaptic function [32,45,46]. As LTP and LTD modify the cellular proteome through the simultaneous recruitment of protein synthesis and degradation; it stands to reason that homeostatic scaling mechanisms may also employ a functional synergy of the two to recompense for the changes brought about by unconstrained Hebbian processes.

AMPAR-mediated currents decrease more than NMDAR currents during chronic network hyperactivity [14,18] and unlike NMDARs, the turnover of AMPARs is translation-dependent [47]. The reduced level of sAMPARs following chronic hyperactivity is reset to basal level by the co-application of protein synthesis and proteasome inhibitors, suggesting that the combined action of translation and degradation affects post-synaptic scaling specifically through sAMPARs. Similar to the observations in synaptic upscaling [13], restricting changes to the sAMPAR abundance by the inhibition of GluA2-endocytosis using GluA2_3Y_ peptide also blocks synaptic downscaling (Figure 2); reinforcing that AMPARs indeed remain the end-point effectors despite changes to the proteome.

### Association of the translation and degradation apparatus is RNA-dependent

The co-localization of polyribosomes and proteasomes in neuronal sub-compartments suggests that for translation and proteasomal degradation to work in tandem, physical proximity between the two modules cannot be ruled out [30,31]. Polysome analysis showed the co-sedimentation of members of the 19S proteasome (Rpt1, Rpt3 and Rpt6 subunits) and the 20S proteasome (α7 subunit) along with translation initiation factors such as eIF4E and p70S6 kinase, a downstream effector of mTORC1. Abrogation of the sedimentation pattern in the presence of RNase or EDTA, is indicative of an RNA-dependent direct interaction between translation and protein degradation (Figure 3). RNAse treatment also abolished the direct interaction between proteasome subunits and translation regulators within the polysome, suggesting that actively-translating transcripts act as a scaffold to link the translation and proteasome machineries (Figure 4). Such existence of direct interaction between polyribosomes and catalytically active proteasomes allows close temporal coordination between translation and protein degradation.

Does chronic hyperactivity influence the association of proteasome and translation regulators within the polysome? The bicuculline-induced enrichment of 26S proteasome, phosphorylated S6, p70 S6 kinase and its phosphorylated form, and eEF2 in polysomes is indicative of activity-dependent proximity between the translation and degradation machineries (Figure 5). Interestingly, eEF2 has been previously characterized as a biochemical sensor for synaptic scaling [17]. Trim32, an E3 ligase, was enriched in polysomes whereas MOV10 and Argonaute to be depleted from polysomes on chronic bicuculline treatment. Trim32 and MOV10 were also present in polysomes in basal conditions (Figure 5).

Previously, MOV10 and Trim32 have been implicated in miRISC-independent functions to modulate RNA modification [48], stability [49] and transcription [50] respectively. Density gradient fractionations of cytoplasmic lysates obtained from non-neuronal systems have revealed that Argonaute cosedimented with miRNAs in polysome fractions [51,52]. MOV10 was also found associated with polysomes [24]. Interestingly, Trim32 and Argonaute co-immunoprecipitated with MOV10 from neurons suggesting that they closely interact with each other and are members of the miRISC (Figure 4F). Apart from this observation, polysome association of Trim32, MOV10 and Argonaute is bicuculline responsive (Figure 5). We found that chronic bicuculline stimulation triggers a change in the association of Trim32 and MOV10 with Argonaute (Figure 6). Evidence of their direct physical interaction led us to infer that both MOV10 and Trim32 are part of the miRISC. Hence, association of Argonaute, Trim32 and MOV10 with polysome can be representative of the association of the miRISC with polysomes; at-least in the context of synaptic downscaling.

How does the proteasome remain associated with actively translating mRNAs? We have identified that HspA2 (Hsp70 family), a chaperone protein, remains tethered to proteasomes and polysomes (Figure 4-5). Hsp70 family of proteins is known to influence both the synthesis and degradation of proteins by their association with 26S proteasomal subunits [53], translation initiation factors [54]. Hspa2 has been shown to be an interacting partner of the miRISC [55]. Therefore, HspA2 could potentially function as a proteostasis coordinator which includes members of the proteasome, translation regulators and chaperon proteins.

### Bicuculline mediated regulation of Trim32 and MOV10 causes miRISC remodeling during downscaling

We found that during chronic bicuculline treatment, translation of Trim32 precedes the degradation of MOV10. The alternative possibility that MOV10 degradation leads to increased *de novo* translation of Trim32, is not supported, since protein synthesis inhibition by anisomycin leads to MOV10 rescue (Figure 6). Loss of Trim32 prevented bicuculline-induced polyubiquitination and subsequent degradation of MOV10, suggesting that Trim32 is the only E3 ligase marking MOV10 for degradation during synaptic scaling (Figure 6). Immunoprecipitation of Argonaute from cultured neurons following chronic bicuculline treatment showed MOV10 to be depleted and Trim32 to be enriched, without any change in Dicer expression. Previously, changes to the components of the miRISC have been termed as miRISC remodeling [25] and such compositional changes result in alteration of miRISC activity [23,56]. Our observations therefore emphasize that such compositional changes within the miRISC, or miRISC remodeling, is effectuated through the reciprocal translational-degradation of the Trim32-MOV10 axis. Prolonged bicuculline treatment also did not influence the Argonaute level, indicating that specific components of the silencing complex are targeted during scaling (Figure 6 and S3).

A recent study has demonstrated that a slow turnover of plasticity proteins (measured at 1,3 and 7 days in cultured neurons) is essential to create long-term changes to the neuronal proteome during both up and down-scaling [57]. The authors have argued that the slow turnover rate is more energy-saving and therefore a preferred cellular mechanism. At the same time, this study also identifies a very small fraction of previously reported scaling factors with fast turnover rates specifically influencing up- and down-scaling. Our reports support the latter findings, where we observe that both the increase in Trim32 synthesis and the resultant degradation of MOV10 happen within 24 hours during synaptic downscaling, suggesting a fast turnover. As both MOV10 and Trim32 are part of the miRISC, their fast turnover rates seems plausible, considering that participation of the miRISC is mandatory to relieve the translational repression of several transcripts encoding plasticity proteins and needs to happen rapidly in order to boost changes to the proteome. Although in terms of energy expenditure the coordinated regulation of translation and degradation is expensive, this cellular trade-off may be necessary to trigger the remodeling of a very limited number of master regulators of the neuronal proteome, such as miRISC, during synaptic downscaling.

### mTORC1 dependent Trim32 translation-driven MOV10 degradation is sufficient to cause synaptic downscaling

What post-synaptic signaling cascade triggers Trim32 translation? We find that chronic bicuculline induction triggers the mTORC1-dependent synthesis of Trim32 with the consequent degradation of MOV10 that is a prerequisite for bicuculline-induced downscaling (Figure 7). Although both p70 S6 kinase and 4EBP2 are downstream effectors of mTORC1, we find that the phosphorylation of p70 S6 kinase exclusively drives Trim32 translation.

After investigating the upstream regulators of the Trim32-MOV10 axis, we focused on identifying the downstream factors that lead to loss of surface AMPAR abundance during downscaling. Most studies have determined the influence of single miRNAs in regulating AMPAR distribution during scaling; however, they have been inadequate in providing a holistic view of the miRNA-mediated control of sAMPAR abundance [19–22]. MiRNA function has been shown to be directly co-related with miRISC activity [23,56]. Hence, we explored how miRISC remodeling contributes to synaptic downscaling *via* the regulation of AMPARs. We found that loss of MOV10 function single-handedly accounted for the loss of sGluA1/A2, accompanied by commensurate decrease in mEPSC amplitude under basal conditions, effectively recapitulating the post-synaptic events during downscaling (Figure 8). MOV10 knockdown did not reduce mEPSC amplitude further following bicuculline treatment. This observation indicates that the loss of MOV10 sets the threshold point of synaptic strength and chronic hyperactivity cannot override this set point (Figure 8).

We genetically manipulated MOV10 and Trim32 so that their expression becomes opposite to that observed during scaling, i.e, we overexpressed MOV10 and performed Trim32 RNAi. We found, that MOV10 overexpression led to a partial occlusion of downscaling (Figure 8). Though myc-tagged MOV10 was amenable for degradation by UPS, yet sustained increased levels of the ectopically expressed MOV10 even after bicuculline treatment was sufficient to cause partial impairment of downscaling. Chronic bicuculline treatment post Trim32 RNAi resulted in an increase in mEPSC amplitudes in neurons, which is commensurate with the enhancement of sAMPARs (Figure 9). Thus, Trim32 knockdown caused a partial impairment of downscaling. We also observed that Trim32 RNAi under basal conditions causes a modest but statistically significant increase in mEPSC amplitude (Figure 9). We anticipate that this could be due to enhanced MOV10 expression following Trim32 knockdown under basal condition (Figure 6E). Since the reversal of Trim32 and MOV10 expression levels lead to the partial abolishment of downscaling in neurons upon chronic bicuculline treatment, we infer that the Trim32-MOV10 mediated miRISC remodeling is pivotally positioned to regulate synaptic scaling.

### mTORC1 triggers miRISC remodeling to regulate Arc synthesis during downscaling

Similar to previous observation [38], our study shows that bicuculline-induced hyperactivity enhances Arc protein, a known regulator of AMPAR removal from synapses. Enhanced Arc translation during chronic bicuculline treatment was blocked by rapamycin, suggesting a regulatory role of mTORC1 in its expression. Furthermore, increase in Arc translation (Figure 10) and concomitant reduction of sAMPARs after loss of MOV10 (Figure 8) demonstrates Arc to be a crucial intermediate between MOV10 degradation and synaptic downscaling.

How Arc expression is regulated during chronic hyperactivity? Our data demonstrates that in the context of downscaling, rapamycin-sensitive Arc expression is driven by post-transcriptional control rather than transcriptional regulation. Chronic hyperactivity-dependent enhanced expression of Arc requires the 3’UTR of the transcript containing multiple miRNA binding sites [40], suggesting an involvement of miRISC in scaling. Trim32 knockdown or mTORC1 inhibition prevents the bicuculline-induced increase in Arc reporter expression. We observed that loss of Trim32 alone does not affect the reporter activity, but MOV10 knockdown is sufficient to enhance it (Figure 10). Since MOV10 RNAi mimics the bicuculline-induced reduction of MOV10 during synaptic hyperactivity, and MOV10 regulates miRISC function, we believe that miRISC remodeling is a key determinant of Arc expression. To further enquire whether change of miRISC activity is sufficient to cause increased Arc expression, we performed the RNAi of Dicer, another key component of the miRISC. Similar to MOV10, loss of Dicer has been shown to inhibit miRISC function [56]. Enhanced Arc reporter activity upon Dicer knockdown emphasizes the requirement of miRISC function in regulating Arc (Figure 10).Arc mRNA levels remain unaffected upon chronic hyperactivity in presence or absence of rapamycin as well as knockdown of MOV10 (Figure 10), suggesting that transcription does not play a role in regulating Arc in the context of bicuculline-induced downscaling.

In contrast to chronic hyperactivity driven loss of MOV10, its polyubiquitination and subsequent localized degradation at active synapses has been shown to occur within minutes upon glutamate stimulation of hippocampal neurons in culture or during fear memory formation in amygdala [23,58]. These observations indicate that MOV10 degradation is a common player involved in both Hebbian and homeostatic forms of plasticity. Hebbian plasticity paradigms triggers homeostatic scaling in neurons as a compensatory mechanism [5]; these two opposing forms of plasticity therefore must involve a combination of overlapping and distinct molecular players. Our data demonstrates the requirement of a rapamycin-sensitive, MOV10 degradation-dependent Arc translation in homeostatic scaling that is distinct from the rapamycin-insensitive dendritic translation of Arc occurring during Hebbian plasticity [59]. We speculate that homeostatic and Hebbian plasticity engages distinct signaling pathways that converge at miRISC remodeling.

Though most homeostatic scaling studies including ours used hippocampal neurons in culture to investigate the mechanistic details, the use of this model leaves a lacuna to evaluate how input-specific gene expression control at selective synapses during Hebbian plasticity influences compensatory changes across all synaptic inputs to achieve network homeostasis. Therefore, physiological relevance of homeostatic scaling needs to be studied in association with Hebbian plasticity in order to delineate factors contributing to proteostasis involving cell intrinsic and extrinsic variables within a circuit.

## Author Contributions

S.B and S.S designed the study. S.S., and B.S. performed all experiments. S.S., B.S., and S.B. analyzed the data. J.P.C. provided critical comments on electrophysiology experiments and manuscript. S.V.S.M gave comments on manuscript. S.S. and S.B wrote the manuscript.

## Acknowledgement

We thank Dr. Dipanjan Roy for helpful discussions, Addgene for lentiviral vectors; Ken Kosik for MOV10 shRNA and myc-MOV10 constructs; Clive Bramham for Arc 3’UTR fused luciferase reporter construct; microscope facility of Regional Centre for Biotechnology, India; Rohini Roy for Trim32 RNAi construct; Utsav Mukherjee, Premkumar Palanisamy, Gourav Sharma, Anupam Das and Surajit Chakraborty for technical assistance and animal maintenance; Karthick Ravichandran for helping with polysome fractionation. This work is supported by Ramalingaswami Fellowship (BT/RLF/Re-entry/32/2011) from the Department of Biotechnology, Government of India and National Brain Research Centre core fund to S.B.

Authors declare no conflict of interest

