## Supplementary Information for "Homeostatic scaling is driven by a translation-dependent degradation axis that recruits miRISC remodeling"

§ Correspondence

**Supplementary Figure**

Figure: S1

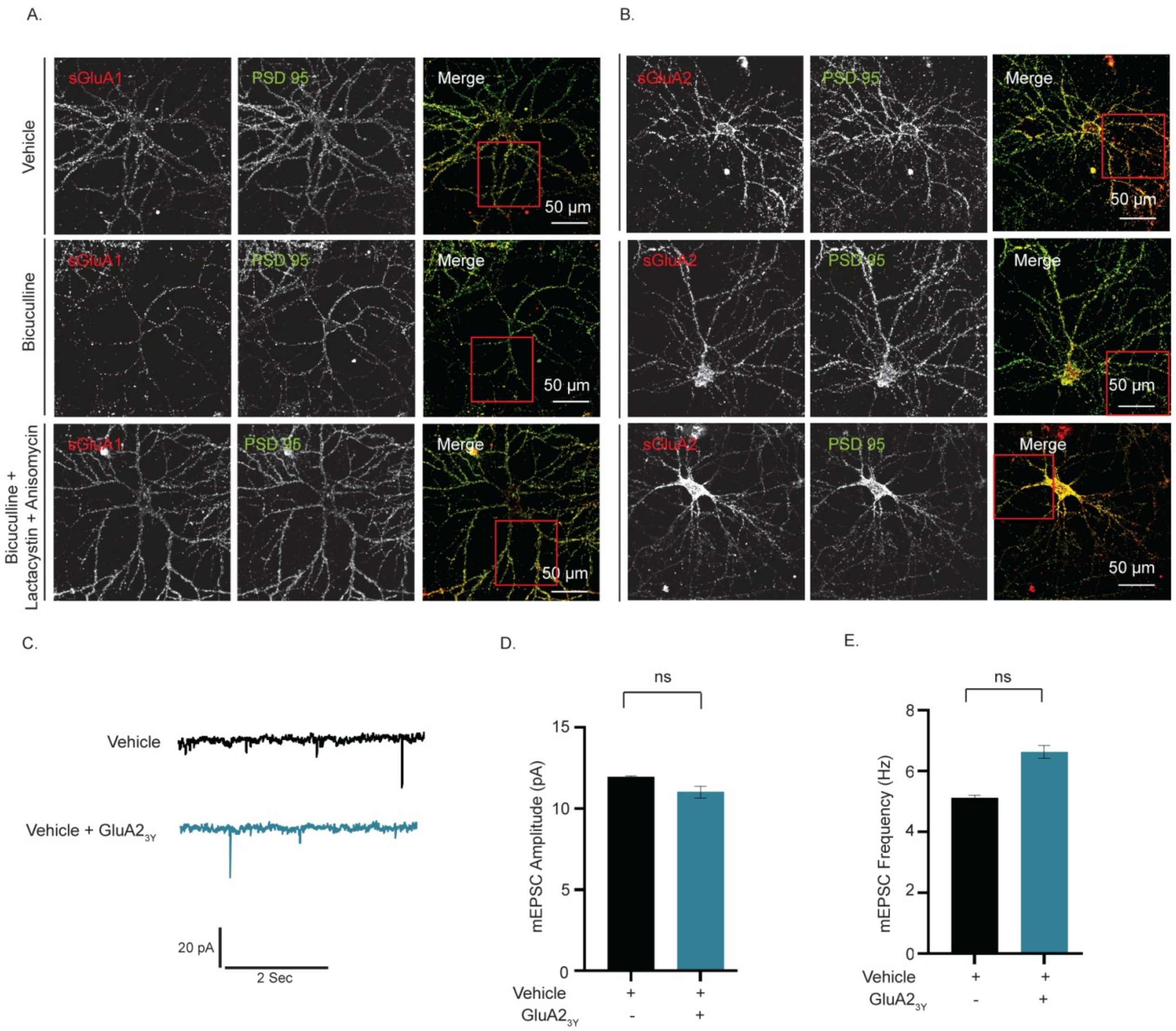

**Supplementary Figure 1: Synaptic downscaling by coordinated control of protein synthesis and degradation involves AMPARs.**

(A-B) Hippocampal neurons were stained for GluA1 (A) or GluA2 (B) and PSD95 as described in Figure 2A-B. Photomicrograph showing images for surface GluA1 or GluA2 (red) and PSD95 (green) and sGluA1/PSD95 or sGluA2/PSD95 (merged). High magnification images of dendrites shown in Figure 2 marked in red square. Scale bar as indicated. Quantitation shown in Figure 2C-D.

(C-E) mEPSCs traces from hippocampal neurons (DIV18-24) treated with vehicle or GluA23y for 24 hours (C) as described in Figure 2E. Scale as indicated. Mean mEPSC amplitudes (D) and frequencies (E) in neurons treated as indicated. n=12. Data shown as Mean ± SEM. One Way ANOVA and Fisher’s LSD.

Figure: S2

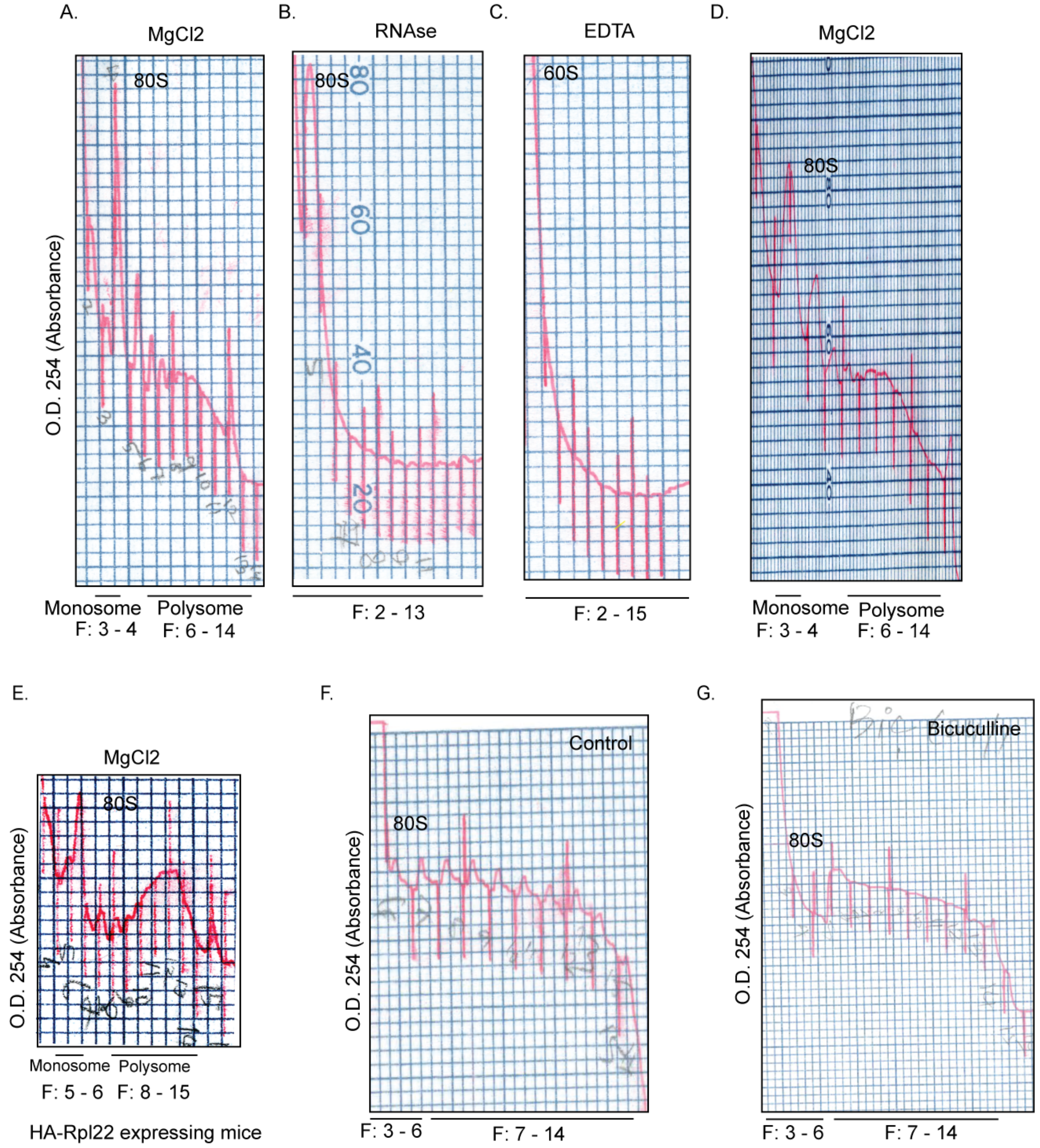

**Supplementary Figure 2: O. $D_{254}$ profile of polysome fractionation**

(A-E) $A_{254}$ profile obtained from spectrophotometer attached gradient fractionator shown in Figure 3 and 4. Traces were drawn from original $A_{254}$ profile obtained from hippocampal cytoplasmic extract treated with $MgCl_2$ (A), RNAse (B), EDTA (C), $MgCl_2$ (D) shown in Figure 3 and $MgCl_2$ treated extract from mouse expressing HA-Rpl22 in excitatory neurons from hippocampus (E) shown in Figure 4. (F-G) $A_{254}$ profile of sucrose density fractions obtained from vehicle (F) or bicuculline (G) –treated cortical neurons. Traces were drawn from these original $A_{254}$ profiles as shown in Figure 5A and 5B.

Figure: S3

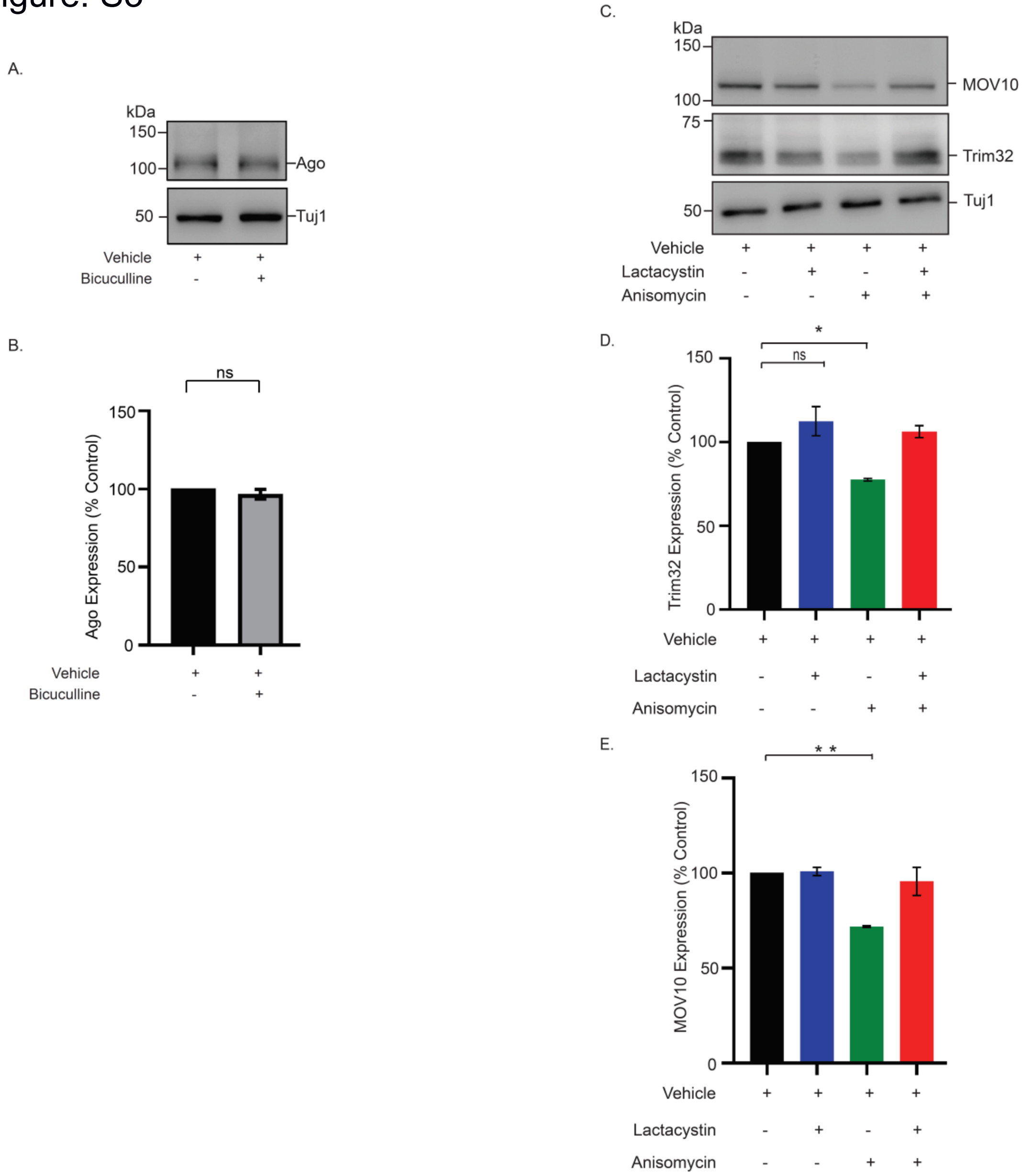

**Supplementary Figure 3: Expression profile of miRISC members under basal condition and bicuculline-induced hyperactivity.**

(A-B). Hippocampal neurons (DIV21) were treated with bicuculline for 24 hours. Photomicrograph showing the expression of Argonaute and Tuj1 as detected by western blot analysis (A) Quantitation of Argonaute expression (B). n=3. ns = not significant. Unpaired 2-tailed t-test.
(C-E) Hippocampal neurons (DIV21) treated with lactacystin, anisomycin and both for 24 hours. Photomicrograph showing the expression of Trim32 and MOV10 as detected by western blot analysis (C). Quantitation of Trim32 (D) and MOV10 (E). Data shown as Mean ± SEM, n=3, *p<0.001 and ** p<0.0003. One Way ANOVA and Fisher's LSD. See also Figure 6.

Figure: S4

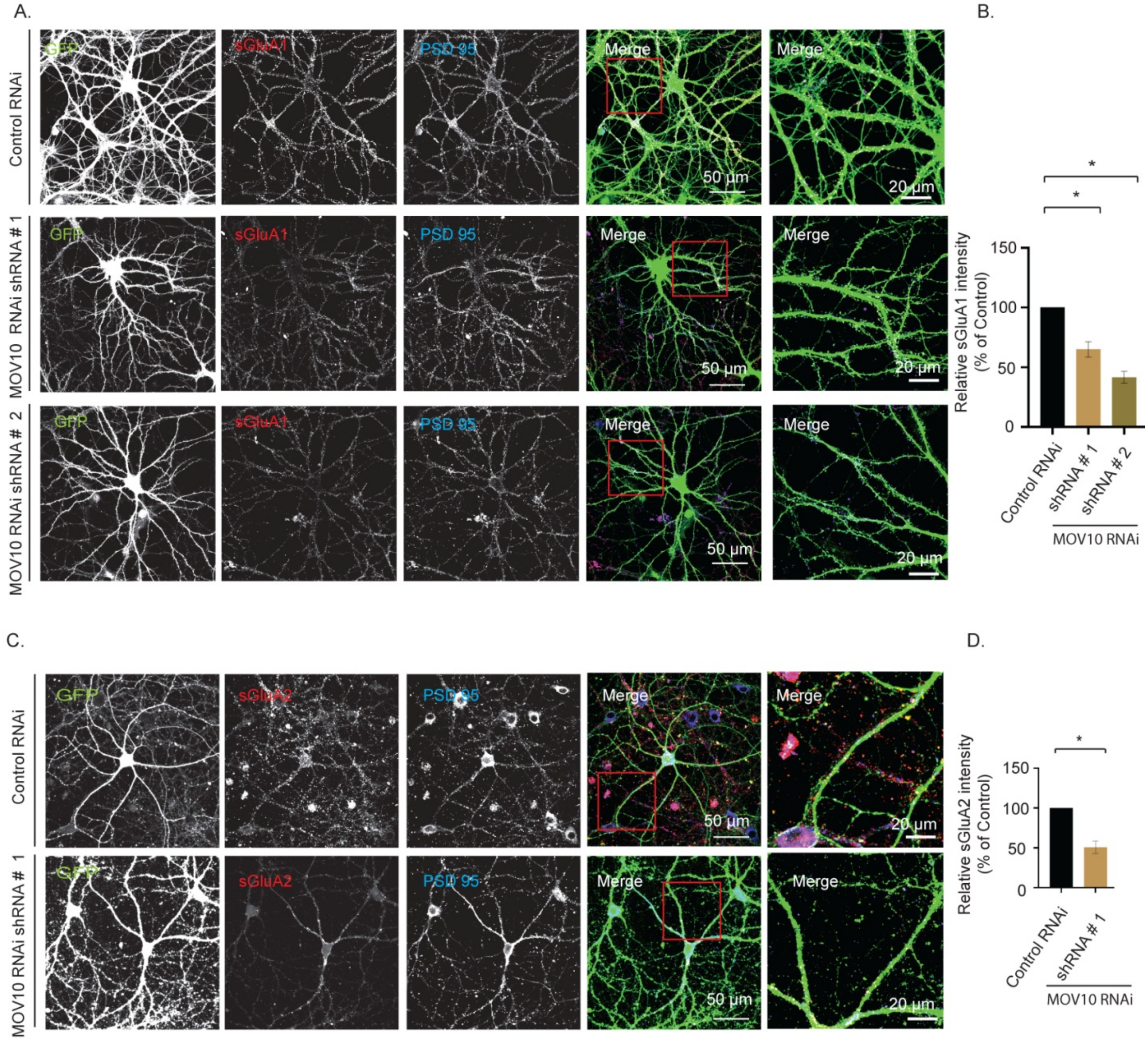

**Supplementary Figure 4: Surface AMPARs expression following MOV10 knockdown**

(A-B) Hippocampal neurons (DIV14-15) transduced with lentivirus expressing two shRNAs against MOV10 (#1 or # 2) along with GFP. Transduced neurons (DIV21-24) were immunostained for surface GluA1 and co-immunostained for PSD95. Photomicrograph showing confocal images of GFP (green), sGluA1 (red), PSD95 (blue) and GFP/sGluA1/PSD95 (merged) (A). High magnification images of dendrites shown in Figure 8 marked in red square. Relative intensity of surface GluA1 particles at the synapse (overlap with PSD95 particles onto GFP expressing dendrites) (B). Normalized intensity of surface GluA1 relative to control was plotted. Data shown as Mean ± SEM. *p<0.01. One Way ANOVA and Fisher's LSD.

(C-D) Hippocampal neurons (DIV14-15) transduced with lentivirus expressing shRNA against MOV10 (#1) along with GFP. Transduced neurons (DIV21-24) were immunostained for surface GluA2 and PSD95. Photomicrograph showing confocal images of GFP (green), sGluA2 (red), PSD95 (blue) and GFP/sGluA2/PSD95 (merged). High magnification images of dendrites shown in Figure 8 marked in red square. Scale as indicated. Relative intensity of surface GluA2 particles at the synapse (overlap with PSD95 particles onto GFP expressing dendrites). Normalized intensity of surface GluA2 relative to control was plotted. Data shown as Mean ± SEM. *p<0.01. One Way ANOVA and Fisher's LSD. See also Figure 8.

Figure: S5A

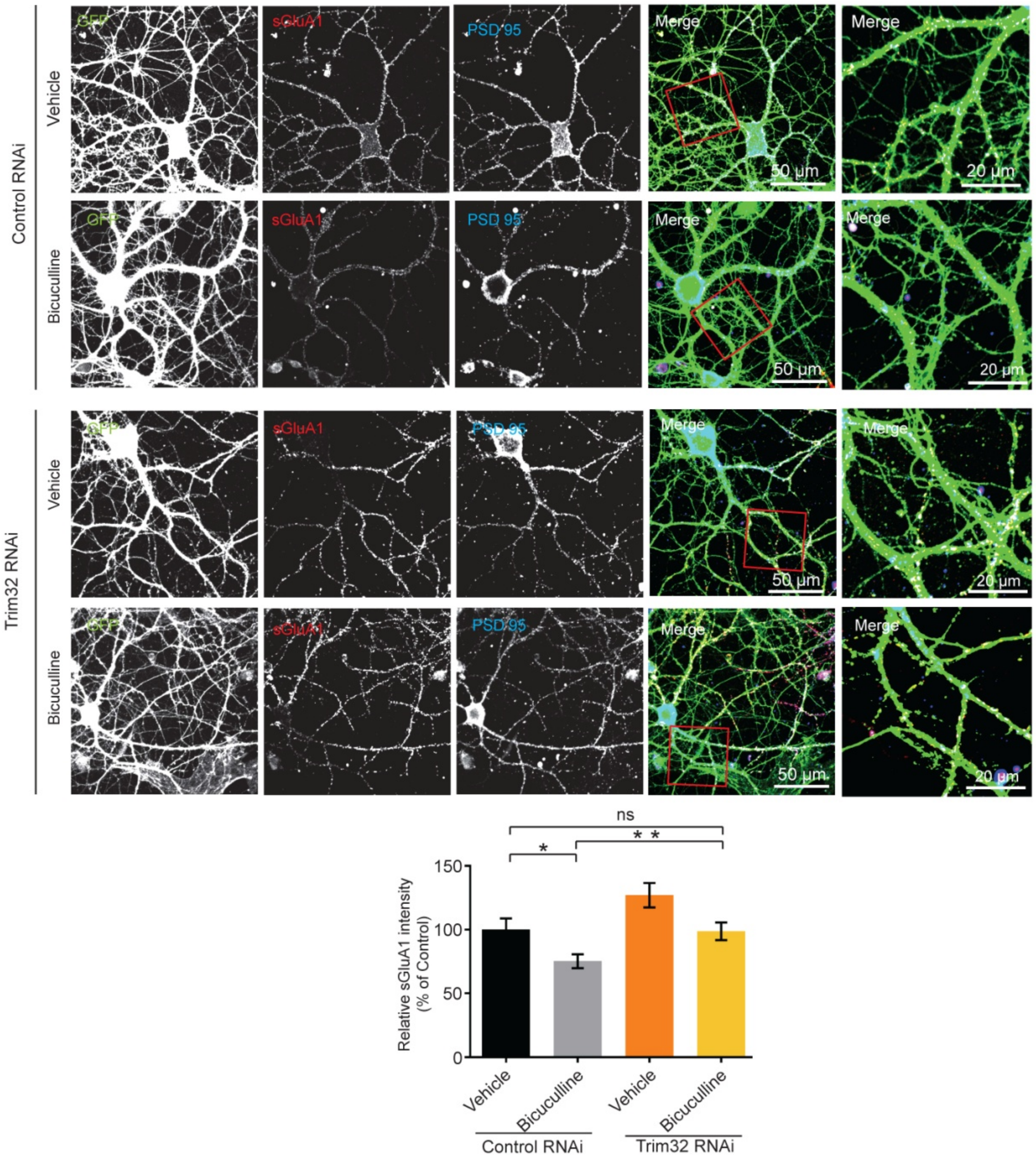

Figure: S5B

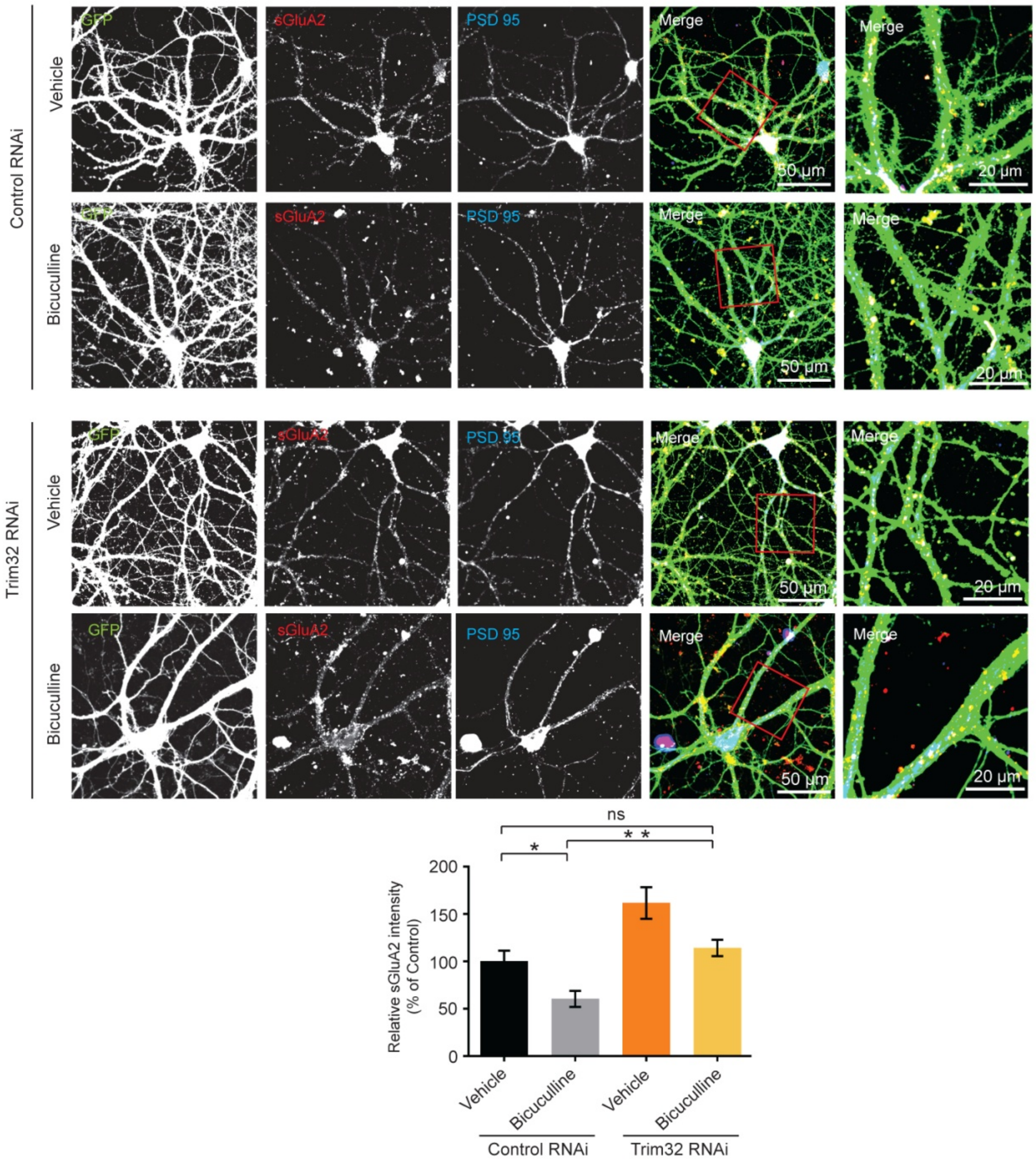

**Supplementary Figure 5: Surface AMPARs expression in bicuculline-induced neurons following Trim32 knockdown**

(A-B) Hippocampal neurons (DIV14-15) transduced with lentivirus expressing shRNA against Trim32 along with GFP. Transduced neurons (DIV21-24) were stimulated with bicuculline for 24 hours and immunostained for surface GluA1 (A) or GluA2 (B) and co-immunostained for PSD95. Photomicrograph showing confocal images of GFP (green), sGluA1/sGluA2 (red), PSD95 (blue) and GFP / sGluA1 or sGluA2 / PSD95 (merged). High magnification images of dendrites shown in Figure 9D – 9I marked in red square. Relative intensity of surface GluA1 (A) or surface GluA2 (B) particles at the synapse (overlap with PSD95 particles onto GFP expressing dendrites). Normalized intensity of surface GluA1 / GluA2 relative to control was plotted. Data shown as Mean ± SEM. *$p<0.02$, **$p<0.0001$ for sGluA1. *$p<0.006$, **$p<0.0005$ for sGluA2. One Way ANOVA and Fisher's LSD. See also Figure 9.

Figure: S6

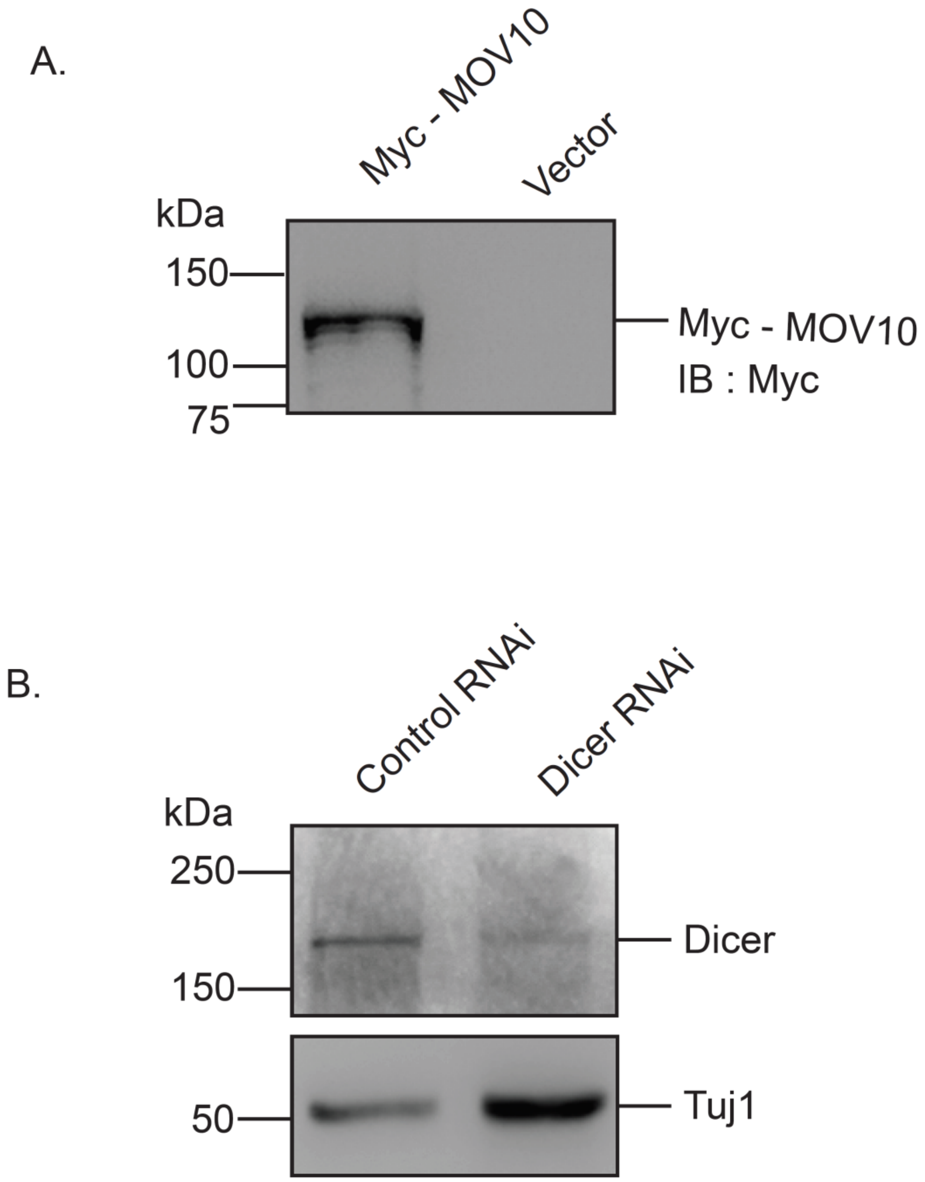

**Supplementary Figure 6: Overexpression of MOV10 and knockdown of Dicer**

(A). Myc-tag MOV10 was transfected in hippocampal neurons (DIV15) as detected by western blot analysis (DIV21) using antibody against Myc. See also Figure 8.

(B) Hippocampal neurons (DIV14) were transduced with lentivirus expressing shRNA against Dicer or control shRNA. Photomicrograph showing effective knockdown of Dicer (DIV24) as detected by western blot analysis using antibody against Dicer. See also Figure 10.

### Supplementary Methods

#### Primary neuronal culture

Hippocampal neuronal cultures from rat (Sprague-Dawley) were prepared and maintained as described previously (Kaech & Banker, 2006). Briefly, hippocampi from embryonic day 18 (E18) pups were dissected, treated with trypsin (0.25%), dissociated by trituration to make single cell suspension and plated onto poly-L-lysine (1mg/mL) coated glass coverslip (160 – 250 cells / $mm^2$). 160 - 170 cells /$mm^2$ were used for electrophysiology and surface labeling experiments. 200 - 250 cells /$mm^2$ cells were used for all biochemical experiments. Cortical neuronal cultures from rat were prepared following previous protocol (Banerjee et al 2009). For polysome experiments, 450-475 cortical cells /$mm^2$ were plated onto 90 mm dishes pre-coated with 1mg/mL poly-L-Lysine (Sigma). Neurons were maintained in Neurobasal medium (Gibco) containing B27 supplements (Gibco) at 5% $CO_2$ / 37°C up to 22-25 days prior to commencement of experiments. Animal experiments were performed with the approval of the Institutional Animal Ethics (IAEC) committee of National Brain Research Centre.

#### Pharmacological Inhibitors:

Primary hippocampal neurons aged DIV 21-24 were treated with bicuculline (10μM,Tocris ), lactacystin (10μM, AM systems) and anisomycin (40 μM, Sigma ) alone or in combination for 24 hours before further analysis. For mTORC1 inhibition experiments, LY2584702 Tosylate, a competitive inhibitor of p70 S6 Kinase was added (final concentration 2μM) to cultured hippocampal neurons for 24 hours.

#### Lentivirus production and transduction

Lentivirus preparations and transduction into hippocampal neuronal cultures were performed as described previously (Banerjee *et al*, 2009). Validated shRNA against Trim32 (TATACCTTGCCTGAAGATC) (Schwamborn *et al*, 2009) or Dicer 1 (GCATGGTGGTGTCGATATT) (Kumar *et al*, 2007) was cloned into MluI and ClaI sites of pLVTHM vector (Addgene) and verified by sequencing. pLVTHM vectors containing MOV10 shRNA cassettes (sh#1:TTATACAAGGAGTTGTAGGTG) or (sh#2: ACTTAGCTCTAGTTCATAACC) (Banerjee *et al*, 2009) and non-targetting control (ATCTCGCTTGGGCGAGAGTAAG) were used for lentivirus preparation. Lentivirus particles were produced by co-transfection of 20μg transfer vector (EGFP cassette under EF1α promoter and shRNA cassette against MOV10 or Trim32 or non-targeting control under H1 promoter in pLVTHM plasmid), 15μg psPAX2 and 6μg pMD2.G into HEK293T cells. The cells were grown in low glucose DMEM media (Gibco) with 10% Fetal Bovine Serum (Gibco) and maintained at 5% $CO_2$ / 37°C. HEK293T ($2\times10^6$ cells) were transfected by calcium phosphate method. Culture supernatant containing lentivirus particles were collected 72 hours post-transfection and concentrated virus stock was prepared by ultracentifugation and viral titres determined

To perform RNAi, hippocampal neurons at *Days in Vitro* (DIV) 14-15 were infected with lentivirus expressing shRNAs against MOV10, Trim32, Dicer and non-targeting control as mentioned. Viral infections were performed at MOI of 1-2 for 6 hours and following infection lentivirus containing media was replaced with fresh Neurobasal media with B27 supplements. Transduced neurons were incubated up to DIV 23-25 with bicuculline (where mentioned) prior to surface labelling and biochemical experiments. Viral infected neurons were tracked by EGFP expression for electrophysiology and imaging experiments.

**Surface labeling of GluA1/A2**

Surface expression of AMPAR subunits (GluA1 or GluA2) was analyzed by live-labeling of hippocampal neurons with primary antibodies against surface epitopes of GluA1 (Millipore) or GluA2 (Millipore), under different conditions. Neurons (DIV 21-24) were immunostained as described previously (Schwarz *et al*, 2010). Prior to immunostaining, neurons were transduced with lentivirus for protein knockdown or treated with vehicle (DMSO), bicuculline (10µM) alone or in combination with lactacystin (10µM) and anisomycin (40µM) for 24 hours. Live neurons were incubated for 15 minutes at 5% $CO_2$ / 37°C with N-terminus specific mouse GluA1 (1:25) or mouse GluA2 (1:10) antibodies diluted in Neurobasal media containing B27 supplements. Following incubation, the cells were washed twice with phosphate buffered saline containing $Mg^{2+}$ and $Ca^{2+}$ (PBS-MC; 137mM NaCl, 2.7 mM KCl, 10 mM $Na_2HPO_4$, 2mM $KH_2PO_4$, 1 mM $Mg_2Cl_2$ and 0.1 mM $CaCl_2$). Cells were then fixed in PBS-MC containing 2% paraformaldehyde and 2% sucrose for 20 minutes at 37°C, washed three times in PBS-MC at room temperature and blocked with PBS-MC containing 2% BSA for 30 minutes at room temperature. Cells were incubated with Alexa-546 conjugated goat-anti-mouse secondary antibody (1:200, Invitrogen) at room temperature for 60 minutes in blocking solution. Cells were permeabilized with PBS-MC containing 0.1% Triton-X-100 at room temperature for 5 minutes. Cells were further incubated with blocking solution for 60 minutes and then with goat PSD95 antibody (1:200, Abcam) for 8 hours at 4°C. Cells were incubated with Alexa-633 or Alexa-488 conjugated donkey-anti-goat secondary antibody (1:200, Invitrogen) at room temperature for 90 minutes. Cells were washed three times with PBS-MC at room temperature and mounted on Vectashield mounting media with DAPI (Vector Laboratories).

**Confocal Imaging and Image Analysis:**

Surface AMPAR subunits on hippocampal neurons following MOV10 knockdown or bicuculline treatment were imaged using a Leica TCS SP8 point scanning confocal microscope with a Leica Plan Apochromat 63X NA = 1.4 oil immersion objective at 1024 × 1024 pixel resolution. GFP and Alexa 488 were excited by 488 nM Argon laser. Alexa 546 and Alexa 633 were excited by solid state and Helium-Neon lasers respectively. GFP, Alexa 488 and Alexa 546 signals were detected by hybrid detectors and Alexa 633 was detected by PMT. All images (8 bit) were acquired with identical settings for laser power, detector gain and pinhole diameter for each experiment and between experiments.

Surface AMPARS for neurons with Trim32 knockdown in the presence or absence of bicuculline were imaged using a Nikon A1 HD25 point scanning confocal microscope with a Nikon Plan Apochromat 100X NA = 1.4 oil immersion objective at 1024 × 1024 pixel resolution. High magnification images were captured using 2X optical zoom. We obtained 4-6 optical sections with 0.5µM step size. GFP were excited by 488 nm solid-state laser. Alexa 546 and Alexa 633 were excited by solid state lasers. GFP and Alexa 546 were detected by GaAsP detectors. Alexa 633 was detected by PMT. All images (16 bit) were acquired under identical conditions of laser power, detector gain and pinhole diameter throughout.

High magnification images, captured from confocal microscopy, were analyzed to observe the intensity of GluA1/A2 expression colocalizing with PSD95 (and GFP for MOV10 RNAi experiments). Images from the different channels were stacked and projected at maximum intensity using ImageJ (NIH). These images were then analyzed using custom written Matlab (Mathworks) programs. First, PSD95 and GFP image signals were thresholded to identify the pixels expressing PSD95 and GFP. Then, the pixels of GluA1/A2, colocalizing with PSD-95 and/or

GFP were filtered and the average global intensity of these colocalizing GluA1 pixels were collected, plotted and further analyzed for statistics.

**Polysome fractionation and TCA precipitation of polysome fractions:**

*Polysome fractionation from rat hippocampus*:
Polysomes from the hippocampi of 8-10 week old SD rats were analyzed following previous protocol (Stefani *et al*, 2004). Following decapitation, the brains were removed and placed in ice-cold HEPES HBSS (1× Hank's basal salt solution, 2.5 mM HEPES-KOH pH 7.4, 35 mM glucose, and 4 mM NaHCO3) containing 100 µg/mL of cycloheximide. From this point on, all experimental steps were done at 4°C. Hippocampi were dissected, pooled and homogenised in homogenization buffer (10 mM HEPES-KOH pH 7.4, 150 mM KCl, 5 mM $MgCl_2$, and 0.5 mM DTT) containing EDTA-free complete protease inhibitors. 1.2mL of homogenization buffer per four hippocampi was used. Tissues were homogenised manually with a Dounce homogeniser and the homogenate was spun at 2000 × g, 10 min at 4°C to discard nucleus. The supernatant (S1) was collected and NP-40 was added to a final concentration of 1% v/v. After 5 min of incubation on ice, S1 was spun at 20,000×g for 10 minutes, the resultant supernatant (S2) was loaded onto a 20-50% w/w linear sucrose density gradient (Sucrose buffer: 10 mM HEPES-KOH pH 7.4, 150 mM KCl, 5 mM $MgCl_2$). In the indicated conditions, EDTA (30mM) or a combination of RNase T1 (Ambion,1000U/mL) and RNase A (Ambion,40U/mL), was added to S2 and incubated for 10mins at room temperature before loading it onto the gradient. The gradients were centrifuged at 40,000 g, 2 hr at 4°C in a Beckman Instruments (Fullerton, CA) SW 41 rotor. Fractions of 0.75 mL volume were collected with continuous monitoring at 254 nm using an ISCO UA-6 UV detector.

*Polysome from HA-Rpl22 mice*:
For polysome from the hippocampi of 8-10 week old HA-Rpl22 transgenic mice, exact protocols as above were followed. Where indicated, tissue lysates were treated with a combination of RNAse T1 (Ambion, 1000U/mL) and RNAse A (40U/mL) prior to loading onto the linear density sucrose gradient.

*Polysome fractionation from rat cortical cultures:*
Cortical cultures at DIV 21 were incubated with 10uM bicuculline or vehicle for 24 hours. Polysomes were prepared from cortical cultures following a previous protocol (Stefani *et al*, 2004) with minor modifications. Seven 90 mm dishes plated with cortical neurons was used in each case. Post-incubation, cells were harvested in ice-cold HEPES-HBSS containing 100ug/mL cycloheximide. Following centrifugation at 3220×g, cells were lysed using homogenization buffer (20 mM HEPES-KOH pH 7.4, 5 mM $MgCl_2$, 150 mM KCl, 0.5 mM DTT) containing 100ug/mL cycloheximide, EDTA-free complete protease inhibitor (Roche, 1 tablet per 5 mL) and 40U/mL RNAse inhibitor (Roche) . All steps were carried out at 4°C. NP-40 was then added to a final concentration of 0.3% of total lysate volume and lysates incubated for 5 minutes on ice, before centrifugation at 12000×g for 30 minutes at 4°C. Supernatants containing equal amount of protein were loaded onto 20%-50% w/w sucrose density gradient (described above) and centrifuged at 40,000×g, 4°C for 3 hours in SW 41 rotor. The gradients were then fractionated as above.

*TCA precipitation*:
Sodium dodecyl sulphate was added to a final concentration of 0.015% to each polysome fraction and incubated for 30 minutes in room temperature. Tri-chloroacetic acid (TCA) was added to polysome fractions at 25% of their volume. All the fractions were incubated for 30 minutes post

TCA addition followed by centrifugation at 13,000×g for 30 minutes at RT. The pellets were washed with ice-cold acetone (Merck) twice and dried. Acetone residues were allowed to evaporate and the pellets were resuspended in Laemmli buffer.

**Proteasome activity assay**

Proteasome activity in monosome and polysome fractions was analyzed by 20S Proteasome Assay Kit (Enzo Lifesciences) as per manufacturer's protocol. Briefly, 20S proteasome chymotrypsin-like activity was tested by incubating 80µl of each fraction with Suc-LLVY-AMC fluoregenic peptide substrate with or without epoxymycin (500nM) for 15 minutes at 30ºC. Fluorescence was detected by fluoremeter (Tecan).

**Immunoprecipitation experiments:**

**i.** *From HA-Rpl22 mice*:

HA-tagged ribosomes from adult male mice were immunoprecipitated following previous protocol (Sanz *et al*, 2009) with minor modifications. RiboTag mice were crossed with CamKII-Cre mice and CamKII-Cre:RiboTag offspring expressing HA-epitope-tagged-Rpl22 were selected by genotyping. Anti-HA-tagged beads (200µl) were washed twice with citrate-phosphate buffer pH-5, (24mM citric acid, 52mM dibasic sodium phosphate) and allowed to equilibrate twice for 5 minutes each in immunoprecipitation buffer (50mM Tris pH-7.5, 100mM KCl, 12mM $MgCl_2$, 1% Nonidet P-40). Hippocampi from four adult (8-10 week old) HA-Rpl22 male mice were taken for preparing homogenates, along with the same number of age-matched RiboTag mice who do not express epitope-tagged Rpl22. Hippocampi were rapidly removed and weighed before homogenization in (10% w/vol) polysome buffer (50mM Tris pH-7.5, 100mM KCl, 12mM $MgCl_2$, 1% Nonidet P-40(NP-40), 1mM DTT, 100µg/mL cycloheximide, EDTA free Roche Protease inhibitor cocktail, 200U/mL RNAse Inhibitor) using a Dounce homogenizer. Homogenates were then pelleted at 5000g, 10 minutes at 4ºC followed by collection of supernatant and re-centrifugation of the supernatant at 10,000g for 10 minutes at 4ºC to create a post-mitochondrial supernatant. The supernatant was pre-cleared with protein-G agarose beads (Invitrogen) for 1 hour, followed by centrifugation at 8000g, 4ºC, for 10 minutes to remove the beads. The pre-cleared supernatant (250µl) was then incubated with the equilibrated anti-HA tagged affinity matrix for 6 hours with continuous mixing. The matrix was recovered by centrifugation at 8000g, 4ºC for 15 minutes followed by two washes with high salt buffer HS-150 (Tris 50mM pH-7.5, KCl 150mM, $MgCl_2$ 12mM, 1% NP-40, DTT 1mM, 100µg/mL cycloheximide, protease and RNAse inhibitors as above) for 5 minutes and two washes with high salt buffer HS-300 (Tris 50mM pH-7.5, KCl 300mM, $MgCl_2$ 12mM, 1% NP-40, DTT 1mM, 100µg/mL cycloheximide, protease and RNAse inhibitors as above) for 5 minutes. All procedures were done at 4°C. The pellets were boiled in Laemmli buffer and supernatant was used for analysis.

**ii.** *From polysome fractions*:

Sucrose gradient fractionations were performed from the hippocampus of mice expressing HA-Rpl22 (see above). High density sucrose fractions from Fraction # 7 to # 15 were pooled from polysomes of each condition (with or without RNAse) and diluted using homogenization buffer (see above). Immunoprecipitation from the pooled fractions was performed following a previous protocol (Sanz *et al*, 2009) using HA antibody-conjugated agarose beads (Biolegend). Incubation with the HA-tagged antibodies was allowed to take place for 18 hours in order to capture all HA-associated protein complexes from the diluted solution. All procedures performed at 4°C. The immunoprecipitated pellet was washed using a high salt buffer (10mM HEPES-KOH pH 7.4, 200mM KCl, 5mM $MgCl_2$ along with protease inhibitors) and boiled in Laemmli buffer. The supernatant was used for western blot analysis.

**iii.** *From wild-type Sprague-Dawley rats*:

Immunoprecipitation of 26S proteasome subunits and MOV10 were performed from adult Sprague Dawley (SD) rats. Hippocampi of four adult (8-10 week old) male Sprague Dawley rats were collected and homogenized in tissue lysis buffer (50mM Tris pH-7.5, 150mM NaCl, 1% NP-40, 2mM EDTA, Roche protease inhibitor cocktail, 200U/mL Invitrogen RNAse inhibitor, and phosphatase inhibitor cocktail (Sigma)) (10% w/vol) using a Dounce homogenizer. Prior to this, recombinant protein G-agarose beads (Invitrogen) were equilibrated in wash buffer WB-150 (10mM Tris pH8, 150mM NaCl, and 0.1% NP-40) twice for 5 mins each and centrifuged at 5000g for 2 minutes at 4ºC to recover. The homogenates were centrifuged at 2000g, 4ºC for 10 minutes followed by collection of supernatant and re-centrifugation at 10,000g at 4ºC for 15 minutes to get a post-mitochondrial supernatant. Protein content of the supernatant was measured using the BCA protein estimation method (Pierce). 2% of the total protein content was kept aside as total input and the remaining was divided into two parts having equal protein content (~250µl each); one to be used for isotype control and the other for experiment purposes. Protein-G agarose (Sigma) beads pre-blocked with 3% BSA were added (20µg) to each part and allowed to incubate with continuous mixing at $4^0$C for 1 hour. The pre-cleared supernatants were collected by centrifugation at 5000g for 10 minutes at $4^0$C. To the control fraction, 5µg of IgG isotype control was added (Mouse IgG in case of Rpt6 and Rabbit IgG in case of MOV10 immunoprecipitation). To the experimental fractions, 5µg of Rpt6 or MOV10 antibody was added and both fractions were allowed to incubate for 4 hours with continuous mixing. 40µg of protein-G agarose beads were added to the fractions and further incubated for 2 hours. The beads were recovered by centrifugation and washed twice with wash buffer IPP-150 (50mM Tris pH7.5, 150mM NaCl, 12mM $MgCl_2$, 1% NP-40 and 0.5 mMDTT along with RNAse, protease and phosphatase inhibitors, see reagent list) followed by twice with IPP-300 (same constituents as IPP-150 except NaCl concentration is 300mM). In case of Rpt6, a further stringent wash with IPP-450 (450mM NaCl, rest same as IPP-150) was required. All procedures were done at $4^0$C. The total input, control and the immunoprecipitated samples were boiled in Laemmli buffer and stored for further analysis.

**iv.** *From cultured neurons*:

Argonaute was immunoprecipitated from primary hippocampal cultures following previous protocol (Peritz *et al*, 2006). Cultured hippocampal neurons were incubated on DIV 21 with bicuculline (10µM) for 24 hours. The following day, cultures were washed with ice-cold 1X PBS with 5mM $MgCl_2$, and lysed with lysis buffer (20mM Tris-HCl pH 7.45, 200mM NaCl, 2.5mM $MgCl_2$ and 1% NP-40) containing EDTA-free complete protease inhibitor and 100U/mL RNase Inhibitor. Lysates were centrifuged at 16000×g, 20 minutes; to remove cellular debris and supernatants were collected from each condition. 5mg of protein-containing lysate from each condition was used for the IP. 30ul of protein G-agarose (50% slurry, Sigma) equilibrated with immunoprecipitation buffer (20 mM Tris-HCl pH 7.5, 200 mM NaCl, and 2.5 mM $MgCl_2$, 40U/mL RNase Inhibitor and protease inhibitor). Mouse IgG (isotype control) and equilibrated protein G-agarose beads were used to pre-clear the lysates for 2 hours. Pre-cleared lysates were incubated with 10ug of Argonaute protein (Millipore) overnight with continuous perturbation followed by incubation with the BSA-blocked protein G agarose beads (60ul of 50% slurry per mL of lysate) for 3 hours at 4°C. Beads were collected by centrifugation and washed with lysis buffer twice. The beads were resuspended containing 2% SDS and then further incubated with Laemmli buffer at 90°C. Supernatant from each condition was analyzed by western blot.

**Polyubiquitination assay and western blot:**

Rat hippocampal cultures previously transduced with shRNA against Trim32 (Trim32 RNAi) or control shRNA (Control RNAi) were incubated with bicuculline (10 µM) or vehicle for 24 hours on DIV 21. All incubation happened in the presence of lactacystin (10 µM) to capture the spectrum of all polyubiquitinated proteins on bicuculline addition. Following incubation, cells were scraped first with ice-cold phosphate-buffered-saline (1X PBS) containing 5mM $MgCl_2$ and collected by centrifugation at 3220×g at 4°C for 20 minutes. Cells were then lysed with polysome extraction buffer (PEB) (50 mM Tris-HCl pH-7.5, 100mM KCl, 12mM $MgCl_2$, 1mM DTT and 0.1% NP-40) and centrifuged at 16000×g, 4°C for 20 minutes. Protein-G agarose beads were equilibrated with immunoprecipitation (IP) buffer (50 mM Tris-HCl pH 7.5, 100mM KCl, 12mM $MgCl_2$, and 0.05% NP-40). Protein estimation was done and equal amounts of protein for each condition were taken for further analysis. Lysates were pre-cleared for 2 hours using isotype-specific antibody and protein-G agarose beads. Pre-cleared lysates were then incubated overnight with continuous perturbation at 4°C with antibody specific for MOV10 (Bethyl Lab). Protein-G agarose beads (50 µl of 50% slurry) were added to each condition and incubated for further 3 hours. Post incubation, the beads were washed twice with high-salt buffer (50mM Tris-HCl pH 7.5, 300mM KCl,12mM $MgCl_2$ and 1mM DTT), followed by one wash with PEB. The beads were boiled twice in Laemmli buffer for 10 minutes at 90°C and the supernatant collected. The supernatants were run on an 8% denaturing gel and probed with an antibody which recognises only polyubiquitinylated proteins but not monoubiquitinylated ones (FK1 antibody, 1:1000, Enzo).

**Western Blot**:

a) *From immunoprecipitated samples*:

Immunoprecipitated samples were analyzed as per previous protocols (Banerjee *et al*, 2009). Briefly, samples were boiled in Laemmli buffer and equal volumes resolved on 8-10% SDS-PAGE. Post transfer of proteins on nitrocellulose membrane (Millipore), blots were blocked with 5% BSA for 1 hour and probed with primary antibodies overnight. Antibodies used were MOV10 (Bethyl Lab, 1:1000), Trim32 (abcam, 1:250), Dicer (NeuroMab, 1:500), Ago (Millipore, 1:1000), Hspa2 (Millipore, 1:500), Rpt6 (Enzo, 1:500), Rpt1 (Enzo, 1:500), 20S proteasome core (Enzo, 1:500), eEF2 (CST, 1:1000), p70 S6 Kinase (CST, 1:1000), eIF4E (CST, 1:500) and anti-HA (Biolegend, 1:1000). Following extensive washing with Tris-Buffer-Saline containing 0.1% Tween-20 (0.1%TBST), secondary antibody supplied with the CleanBlot HRP detection kit (Thermo Scientific) was used to detect the proteins using standard chemiluminescence detection on X-ray films or the Mini HD9 gel documentation system (UVITEC, Cambridge). Band intensities were quantified by densitometry using ImageJ software.

b) *From polysomes*:

Equal volumes of TCA-precipitated polysome fractions were resolved on 8-15% SDS-PAGE and transferred onto nitrocellulose membranes. Following blocking with 5% BSA, blots were probed with Rpt6, Rpt1, Rpt3, 20S proteasome core and α7 subunit of proteasome (Enzo), eIF4E (CST), p70 S6 kinase and phospho P70 S6 kinase (CST, 1:500), Rp S6 and phospho-Rp S6 (CST, 1:500), Ago (Millipore), Hspa2 (Millipore), Trim32 (abcam) and MOV10 (Bethyl Lab) overnight at 4ºC. Post incubation, blots were washed with 0.1% TBST and probed with appropriate secondary antibodies. Blots were detected using standard chemiluminescence (Millipore) detection on X-ray films or on the Mini HD9 gel-doc. For polysomes obtained from bicuculline-treated cortical cultures, total band intensity (after background subtraction) of all the bands from fractions 7-14 (polysome fractions) were obtained by densitometry on ImageJ and the resultant values normalized with the total area under the curve of all polysome fractions in each polyribosome profile.

c) *From cultured neurons*:

Post incubation with pharmacological inhibitors, cells were washed twice in pre-warmed PBS and collected in Laemmli buffer. Equal volumes of lysates were resolved on 8-10% SDS-PAGE, transferred onto nitrocellulose membrane, blocked with 5% BSA and probed with antibodies against MOV10, Trim32, Ago, Dicer, Arg3.1 (CST, 1:250), p70 S6K and phospho-p70 S6K. Blots were detected using standard ECL chemiluminescence detection (Millipore) and band intensity determined by ImageJ. Blots were normalized to Tuj1. MOV10, Dicer and Trim32 RNAi samples were also detected similarly.

**Electrophysiology**

Whole cell patch clamp experiments were performed using primary hippocampal neurons (DIV18-25). Neurons were incubated with bicuculline (10μM), anisomycin (40μM), lactacystin (10μM), rapamycin (100nM) and GluA23y (10μM) for 24 hours. Neurons were patched with glass micro-electrodes with an open-tip resistance of 3-8MΩ. Cells with series resistance >30MΩ were excluded from the analysis. To measure the excitatory currents, the following composition of internal solution was used: 100mM Cesium gluconate, 0.2mM EGTA, 5mM MgCl2, 2mM ATP, 0.3mM GTP, 40mM HEPES, pH 7.2 (285-290 mOsm). Miniature EPSCs (mEPSCs) were recorded by holding the cells at -70mV in a recording solution consisting of: 119mM NaCl, 5mM Kcl, 2mM CaCl2, 2mM MgCl2, 30mM glucose, 10mM HEPES, pH7.4 (310-320 mOsm) in the presence of 1μM tetrodotoxin and 10μM Bicuculline.

Average of mEPSC events for 300s from each neuron was analyzed and only the events with <-4pA of peak amplitudes, >0.3pA/ms of rise rates, and 1-12ms of decay time constants were selected for the analysis.

All recorded signals were amplified by Multiclamp 700B (Molecular devices), filtered at 10 Khz and digitised at 10-50 KHz. Analog to digital conversion was performed using Digidata 1440A (Molecular Devices). All data were acquired and analysed using pClamp10.5 software (Molecular Devices) and custom Matlab filtering algorithms. Cells with holding currents greater than -100pA were excluded from the analysis, as well as any cell which was unstable during the recording.

**Luciferase assay:**

Hippocampal neurons transduced with MOV10, Trim32 or Dicer shRNAs at DIV 9-10 were transfected at DIV 18 and bicuculline added on DIV 20. Luciferase assays were performed on DIV-21. Cells were co-transfected with Gaussia luciferase (Gluc) containing the complete Arc 3' UTR from mouse brain cDNA and Firefly luciferase (Fluc) cloned in pMIR-Report (Ambion). Luciferase activities of the two reporters were measured using the Dual Luciferase Reporter Assay System (Promega) according to the manufacturer's instructions. Gluc activity was normalized by Fluc.

**qRT-PCR analysis of Arc mRNA:**

Total RNA was isolated from cultured neurons either subjected to MOV10 RNAi or treated with bicuculline and/or rapamycin for 24 hours using Trizol (Invitrogen cDNA was synthesised using SuperScript® III First Strand Synthesis System (Life technologies, Invitrogen).

Following primers were used for qRT-PCR : Forward: 5'-GGGTGGCTCTGAAGAATATT-3',
Reverse: 5'-TGTACTGCAGAAACTCCTTC-3'.qRT-PCR results analyzed by δδCt method.

**Statistical Analysis:**

Statistical Analyses were performed for all experiments. Whole cell patch clamp amplitudes and frequencies were analyzed using one-way ANOVA with post-hoc Fisher's LSD test to test pairwise differences across the groups. Imaging and western blot data were analyzed for statistical significance using one-way ANOVA with post-hoc Fisher's LSD test. Western blot data related to RNAi experiment was analyzed using unpaired t-test with Welch's correction. Data is reported as absolute differences in mean ± SEM for electrophysiology data or percent differences in mean ± SEM for imaging and western blot data between groups.

**KEY RESOURCES TABLE**

| *REAGENT or* | *SOURCE* | *IDENTIFIER* |
|---|---|---|
| *Antibodies* | | |
| Anti-GluR1-NT (N-terminus) Antibody, clone RH95 | Merck Millipore | Cat#MAB 2263;<br>RRID: AB_1977459 |
| Anti-GluR2 Antibody, clone 14C12.2 | Merck Millipore | Cat#MABN1189;<br>RRID: AB_2737079 |
| Anti-PSD95 antibody | Abcam | Cat#ab12093;<br>RRID: AB_298846 |
| eIF4E (C46H6) Rabbit mAb | Cell Signaling Technology | Cat#2067;<br>RRID: AB_2097675 |
| p70 S6 Kinase (49D7) Rabbit mAb | Cell Signaling Technology | Cat#2708 |
| Proteasome 19S Rpt1/S7 subunit monoclonal antibody (MSS1-104) | Enzo Life Sciences | Cat#BML-PW8825; RRID: AB_10541044 |
| Proteasome 19S ATPase subunit Rpt6 monoclonal antibody (p45-110) | Enzo Life Sciences | Cat#BML-PW9265; RRID: AB_10555017 |
| Proteasome 19S Rpt3/S6b subunit polyclonal antibody | Enzo Life Sciences | Cat#BML-PW8250;<br>RRID: AB_10540650 |

| | | |
|---|---|---|
| Proteasome 20S core subunits polyclonal antibody | Enzo Life Sciences | Cat#BML-PW8155; RRID: AB_2171415 |
| Purified anti-HA.11 Epitope Tag Antibody | BioLegend | Cat#901501; RRID: AB_2565006 |
| Anti-Hsp72 Antibody, clone 3G7 | Merck Millipore | Cat#MABE973; |
| Anti-pan Ago Antibody, clone 2A8 | Merck Millipore | Cat#MABE56; AB_10807962 |
| eEF2 antibody | Cell Signaling Technology | Cat#2332; RRID: AB_10693546 |
| Phospho-p70 S6 Kinase (Thr389) (1A5) | Cell Signaling Technology | Cat#9206; RRID: AB_2285392 |
| Anti-TRIM32 antibody | Abcam | Cat#ab96612; RRID: AB_10679378 |
| MOV10 Antibody | Bethyl Lab | Cat#A301-571A; RRID: AB_1040002 |
| Monoclonal Anti-β-Tubulin III (neuronal) antibody (Tuj1) | Sigma Aldrich (Merck) | Cat#T8578; RRID: AB_1841228 |
| GAPDH | Sigma Aldrich (Merck) | Cat#G9545; RRID: AB_796208 |
| Anti-HA.11 Epitope Tag Affinity Matrix | BioLegend | Cat#900801; RRID: AB_2564999 |
| Donkey anti-Goat IgG (H+L) Secondary Antibody, Alexa Fluor 488 | Thermo Scientific (Invitrogen) | Cat#A11055; RRID: AB_2534102 |
| Donkey anti-Goat IgG (H+L) Secondary Antibody, Alexa Fluor 633 | Thermo Scientific (Invitrogen) | Cat#A21082; RRID: AB_141493 |

| | | |
|---|---|---|
| Goat anti-Mouse IgG (H+L) Secondary Antibody, Alexa Fluor 546 | Thermo Scientific (Invitrogen) | Cat#A11030;<br>RRID: AB_144695 |
| Goat anti-Rat IgG (H+L) Secondary Antibody, HRP | Invitrogen | Cat#31470;<br>RRID: AB_228356 |
| Peroxidase AffiniPure Goat Anti-Mouse IgG (H+L) | Jackson ImmunoResearch Inc. | Cat#115-035-003; RRID:AB_10015289 |
| Peroxidase AffiniPure Goat Anti-Rabbit IgG (H+L) | Jackson ImmunoResearch Inc. | Cat#111-035-003;<br>RRID: AB_2313567 |
| AffiniPure Goat Anti-Mouse IgG (H+L) | Jackson ImmunoResearch Inc. | Cat#115-005-062;<br>RRID:AB_2338452 |
| Rabbit IgG Isotype Control | Invitrogen | Cat#10500C; RRID: AB_2532981 |
| FK1 antibody | Enzo LifeSciences | Cat# BML-PW8805 |
| 20S Proteasome core subunits | Enzo LifeSciences | Cat# BML-PW8155 |
| Anti-pan Ago Antibody, clone 2A8 | Millipore | Cat# MABE56 |
| Anti- Dicer clone N167/7 | Neuromab (now sold by antibodies inc) | Cat# 75-196<br>(from antibodiesinc) |
| *Chemicals, Peptides, and Recombinant Proteins* | | |
| (-)-Bicuculline methochloride | Tocris Bioscience | Cat#0131;<br>CAS#: 53552-05-9 |
| Lactacystin | AG Scientific | Cat#SKU L-1147;<br>CAS#: 133343-34-7 |
| Tetrodotoxin | Abcam PLC | Cat#ab120054;<br>CAS#:4368-28-9 |

| | | |
|---|---|---|
| Glutamate Receptor Endocytosis Inhibitor, GluR23y, YKEGYNVYG | AnaSpec, Inc. | Cat#AS-62547; |
| Anisomycin from *Streptomyces griseolus* | Sigma-Aldrich, Inc. | Cat#A-9789;<br>CAS#: 22862-76-6 |
| Rapamycin from *Streptomyces hygroscopius* | Sigma-Aldrich, Inc. | Cat#R-0395;<br>CAS#: 53123-88-9 |
| Recombinant Protein G Agarose | Invitrogen | Cat# 15920010 |
| EDTA-free Protease Inhibitor Cocktail | Roche (Sigma Aldrich) | Cat#**05892791001** |
| LY2584702 tosylate | Sigma | Cat# SML2892 |
| Cycloheximid e | Sigma Aldrich | Cat# C1988;<br>CAS #66-81-9 |
| SUPERaseIn RNase Inhibitor (20 U/µL) | Invitrogen (Ambion) | Cat# AM2694 |
| Phospahatas e Inhibitor Cocktail 1 | Sigma Aldrich | Cat# P2850 |
| RNase A | Thermo Scientific (Ambion) | Cat# AM2269 |
| Rnase T1 | Thermo Scientific (Ambion) | Cat# AM2283 |
| *Critical Commercial Assays* | | |
| BCA protein assay kit | Thermo Scientific (Pierce) | Cat#23227; |
| Clean-Blot IP Detection Kit (HRP) | Thermo Scientific (Pierce) | Cat#21232; |
| Immobilon western chemiluminisc ent HRP substrate | Millipore | Cat#WBKLS0500 |
| 20S Proteasome Assay Kit | Enzo Life Sciences | Cat# BML-AK740 |

| | | |
|---|---|---|
| Dual luciferase Assay Kit | Promega | Cat# E1910 |
| SuperScript™ III First-Strand Synthesis System | Thermo Fischer | Cat# 18080051 |
| *Experimental Models: Cell Lines* | | |
| HEK293T | ATCC | ATCC #CRL-3216<br>RRID: CVCL_0063 |
| *Experimental Models: Organisms/Strains* | | |
| Sprague Dawley Rats | National Brain Research Centre, India | N/A |
| Ribo Tag transgenic mice in C57BL6/J background | The Jackson Laboratory | 029977 |
| T29-1 CamK2α-Cre transgenic mice in C57BL6/J background | The Jackson Laboratory | 005359 |
| *Recombinant DNA* | | |
| pLVTHM (used for preparing lentivirus) | Addgene | Plasmid #12247<br>RRID:Addgene_12247 |
| psPAX2 | Addgene | Plasmid #12260<br>RRID:Addgene_12260 |
| pMD2.G | Addgene | Plasmid #12259<br>RRID:Addgene_12259 |
| *Software and Algorithms* | | |
| ImageJ | NIH | RRID: SCR_003070;https://imagej.net/ |
| GraphPad Prism8 | GraphPad Software | RRID: SCR_002798; http://www.graphpad.com/ |
| MATLAB | The MathWorks, Inc | RRID: SCR_001622;http://www.mathworks.com/products/matlab/ |
| pCLAMP10.5 | Molecular Devices, LLC | RRID: SCR_011323;http://www.moleculardevices.com/products/software/pclamp.html |
